## Supplementary material for "Solving the *where* problem and quantifying geometric variation in neuroanatomy using generative diffeomorphic mapping": documentation

### GDM

*Release 0.0.1*

Feb 12, 2024

### CONTENTS:


1 Introduction 3

2 Installation 9

3 Coordinate Systems 11

4 File Formats 19

5 Input specification via Transformation Graph Interface 23

6 Output Specification 27

7 Examples 29

8 Function reference 49

9 Work in progress 51

10 Installing 53

11 Examples 55

12 Important Functions 57

13 Module and function documentation 59

14 Web interface 61

Generative diffeomorphic mapping (GDM) is a deformable image registration algorithm designed for aligning multimodal neuroimaging datasets to one another for subsequent analysis. Our package has several important novel features including estimation of any differences in contrast or color between datasets, identification of missing tissues or artifactions, diverse geometries such as mapping 3D volumes to a sequence of 2D datasets, and complex multimodality registration setups described by transformation graphs.

This documentation can automatically be rendered as a pdf using the sphinx package but is best viewed in html (https://twardlab.github.io/emlddmm/build/html/index.html). The pdf may have missing or broken links and images.

#### CHAPTER ONE

### INTRODUCTION

The purpose of our pipeline is to coregister neuroimaging datasets of different modalities and with different coordinate systems. We support 3D to 3D mapping, 3D to 2D mapping (e.g. mapping to serial sections), and 2D to 2D mapping (e.g. rigidly aligning slices with different stains).

We perform registration using diffeomorphisms (with time varying velocity field parameterization) and affine transforms. These transformations can be composed to map data between coordinate spaces and between single specimens and common coordinate systems.

Examples of typical workflows are below. In the diagrams below, each arrow represents the computation of a transformation. By following arrows in the forward or reverse direction, all data can be reconstructed in any of the available spaces. A minor caveat is that only low resolution 2D summary data can be reconstructed in a 3D space.

#### 1.1 Example workflow: STP mapping

A common setting is when we do not have serial section data. For example we may map the Allen atlas to a single 3D STP image. We will need to superimpose atlas labels on the STPT image, and transform the STPT image to match the shape of the atlas.

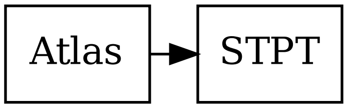


Fig. 1: An example task of 3D to 3D mapping between an atlas and a Serial Two Photon Tomography dataset.

#### 1.2 Example workflow: Alternating sections to atlases

A typical example is to image a mouse brain using serial sections. Alternate sections are stained for Nissl, or for a specific fluorescent tracer. The pipeline will rigidly register fluorescent slices to neighboring Nissl slices, and will deformably register the Allen CCF Nissl atlas onto the 3D stack of Nissl slices. This allows us to map the anatomical labels from the atlas onto our slices. On each slice, we can quantify cell counts or fluorescence in atlas regions. In 3D we can quantify tracer or cell density.

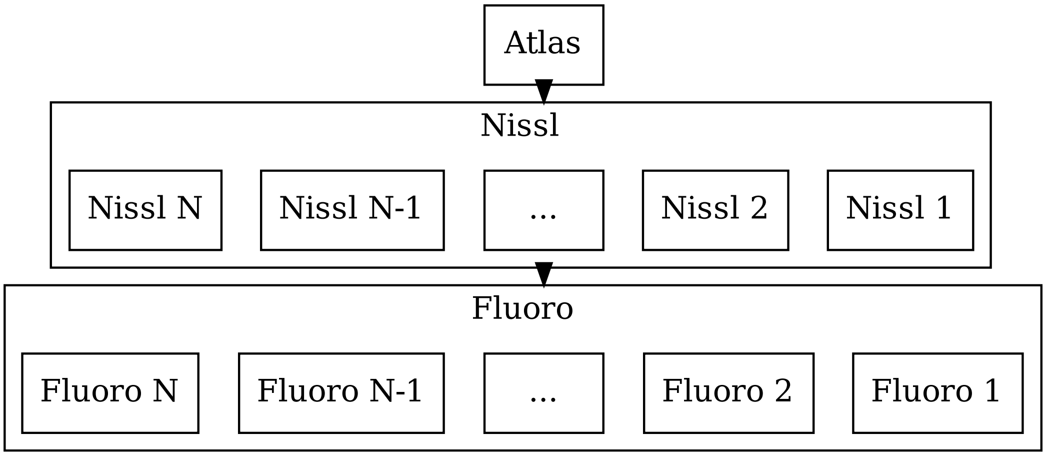


Fig. 2: We map our 3D atlas onto a series of 2D nissl images. We also map our 2D Nissl images to their nearest fluorescent image

Note that any time our pipeline registers a 3D volume to a set of 2D slices, a new space is automatically created called a "registered" space. In this space, all the Nissl sections will be rigidly aligned into a 3D reconstruction.

#### 1.3 Example workflow: Ex vivo MRI

Another example is when MRI is available for a specimen. We typically have an in vivo MRI, ex vivo MRI, and serial section microscopy. The registration tasks are: i) ex vivo to in vivo, ii) ex vivo to serial sections, iii) ex vivo to atlas. We may wish to reconstruct our data in any of the three spaces (in vivo, ex vivo, or atlas). Here the ex vivo MRI plays the role of a common space that is mapped to everything.

Again, a reconstructed space will be automatically created.

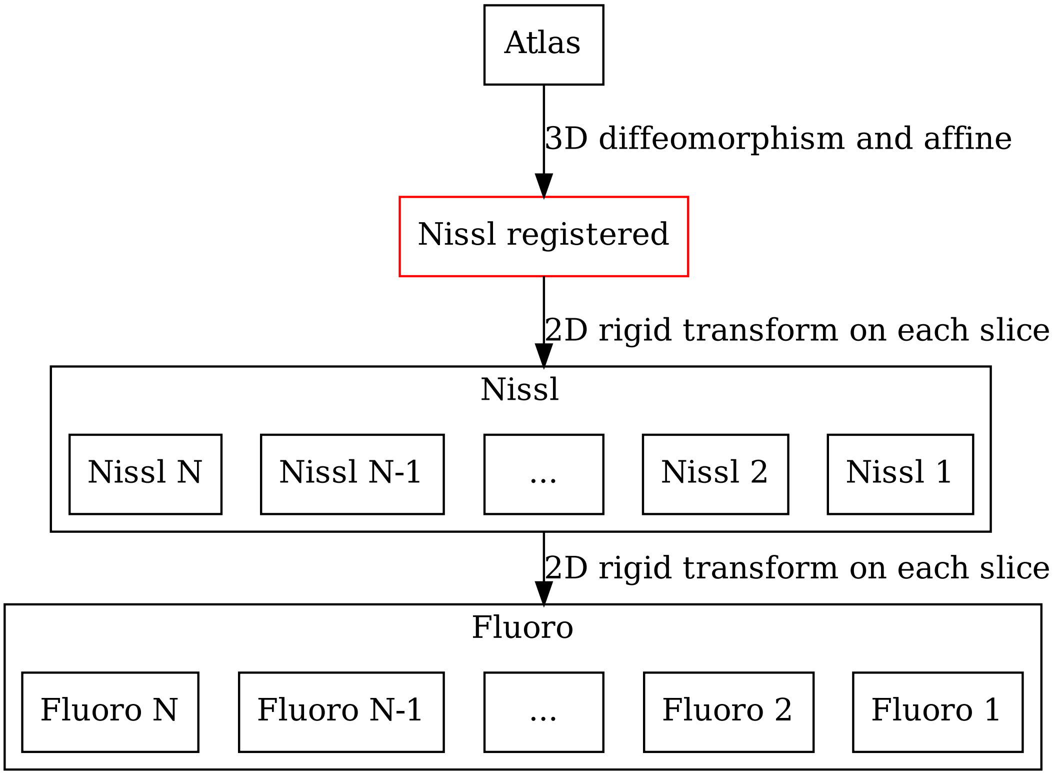


Fig. 3: For any 3D to 2D map, a registered space is automatically created (shown in red). No input data is associated with this space, but images can be reconstructed into this space.

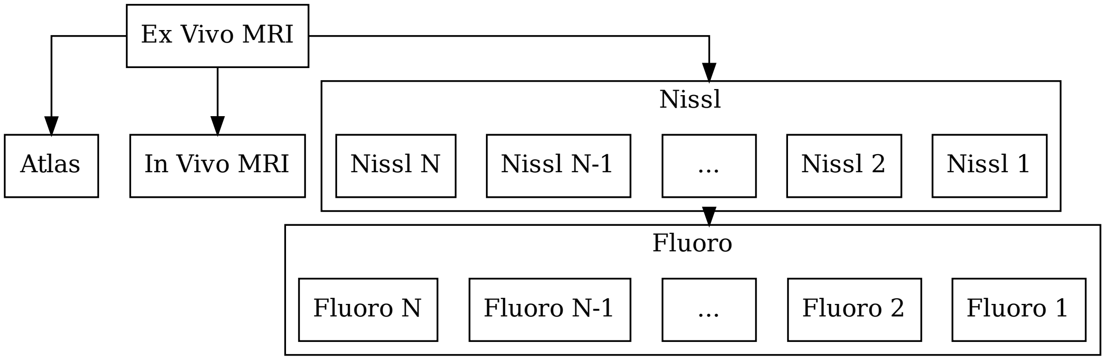


Fig. 4: We may also include in vivo and ex vivo mri.

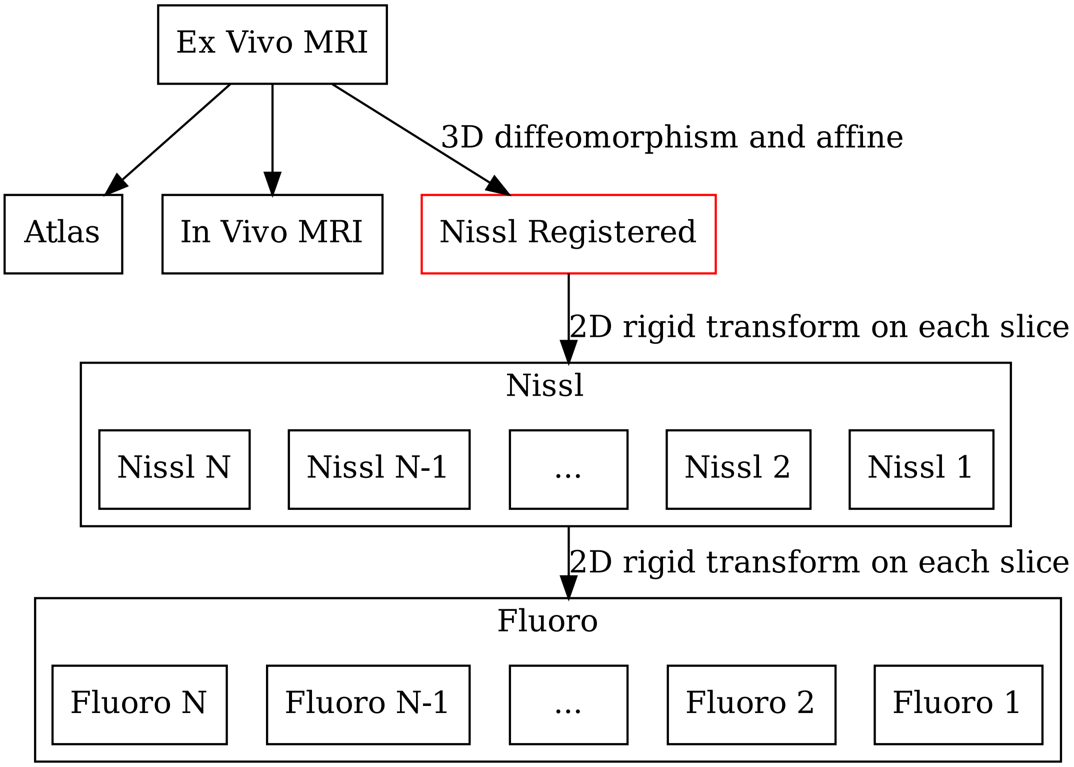


Fig. 5: For any 3D to 2D map, a registered space is automatically created (shown in red). No input data is associated with this space, but images can be reconstructed into this space.

#### 1.4 Example workflow: Arbitrary layout

In general, a registration task can be formulated by a directed acyclic graph. Each node in the graph is a "space", which may have more than one image associated with it. Each arrow in the graph is a registration task.

We have built infrastructure to perform necessary maps, and compose transforms to reconstruct any dataset in any space.

CHAPTER

TWO

### INSTALLATION

Clone the repository:

```
git clone github.com/twardlab/emlddmm
```

Change to the directory emlddmm has been cloned to and install the requirements:

```
pip install -r requirements.txt
```

When running interactively in python, add the appropriate path:

```
import sys
sys.path.append('PATH_TO_EMLDDMM_LIBRARY')
```

When running from command line:

```
python PATH_TO_EMLDDMM_LIBRARY/transformation_graph_v01.py --in INPUT_FILENAME
```

For details on the command line interface, see *input specification*.

### CHAPTER THREE

### COORDINATE SYSTEMS

Our pipeline computes transformations between pairs of spaces. Each space can include one or more images that are sampled on the same voxel grid. Spaces are defined by an origin and an orientation, and often a voxel size. Depending on conventions, these may be specified relative to anatomy in an image (common for atlases) or relative to the sampling grid the image was obtained on. The latter case is common because generally we don't know where the anatomy is until after we have solved a registeration problem.

#### 3.1 Atlas spaces

Coordinate systems associated with several different atlases are described here.

##### 3.1.1 Mouse atlas

We adopt the Allen Institute's Common Coordinate Framework (CCF) atlas, and use its structure annotations, its 3D Nissl image, and its 3D STPT average template image.

Note that the STPT image and annotations are left right symmetric in this data. Nissl images are not left right symmetric.

The CCF NRRD files do not specify a coordinate system correctly. We build our own right handed coordinate system which is described below.

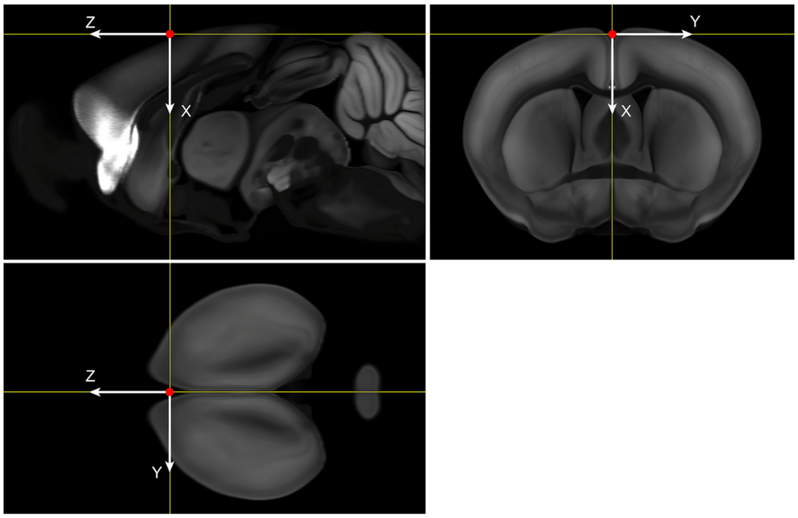

###### Coordinate directions

We chose “x” to point from the brain’s superior to inferior, “y” to point from the brain’s left to right (note this is often displayed by pointing from the right side of the screen to the left, i.e. radiological convention), and “z” to point from posterior to anterior (caudal to rostral).

###### Coordinate origin

We chose “x” to point from the brain’s superior to inferior, “y” to point from the brain’s left to right (note this is often displayed by pointing from the right side of the screen to the left, i.e. radiological convention), and “z” to point from posterior to anterior (caudal to rostral).

We choose the coordinate origin to be our best estimate of the bregma location, based on previous mappings to MR data with skull. Location on Average Template (volume size 8 x 11.4 x 13.2 mm): * x=930 µm from dorsal end of the volume * y=5700 µm from right end of the volume * z=8000 µm from posterior end of the volume

###### Pixel size

This dataset is available in 10, 25, 50, and 100 micron isotropic voxel size. We typically perform registration at 50 microns.

###### Other information

**Warning:** Different versions of annotations are available. Unless otherwise specified, we use CCF version 3. This can be downloaded here: http://download.alleninstitute.org/informatics-archive/current-release/mouse_ccf/annotation/ccf_2017/ Note, as of March 2023, there is a new version of annotations called 2022. But we are not using that version.

**Note:** On CSH server: /nfs/data/main/M32/RegistrationData/ATLAS/annotation_50_bregma_LR.vtk /nfs/data/main/M32/RegistrationData/ATLAS/ara_nissl_50_bregma.vtk /nfs/data/main/M32/RegistrationData/ATLAS/average_template[illegible]

##### 3.1.2 Marmoset atlas

For marmoset we typically use the RIKEN atlas described in Woodward 2018 (https://www.nature.com/articles/sdata20189). An image of our coordinate system convention is shown below.

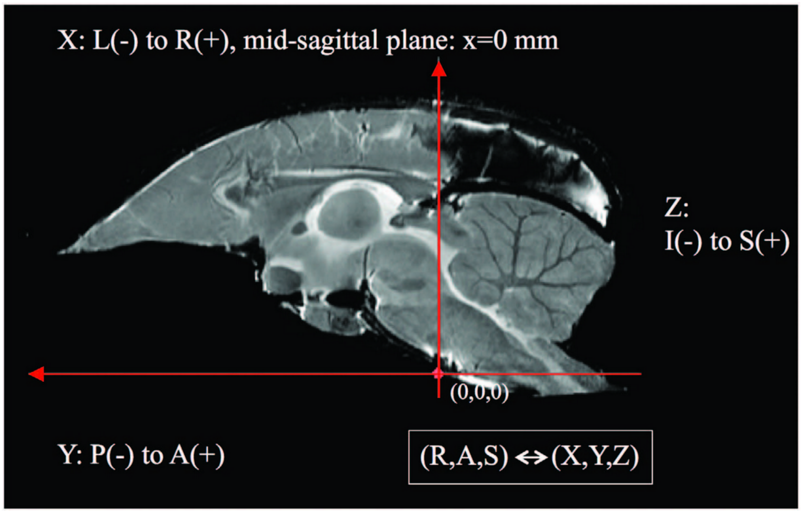


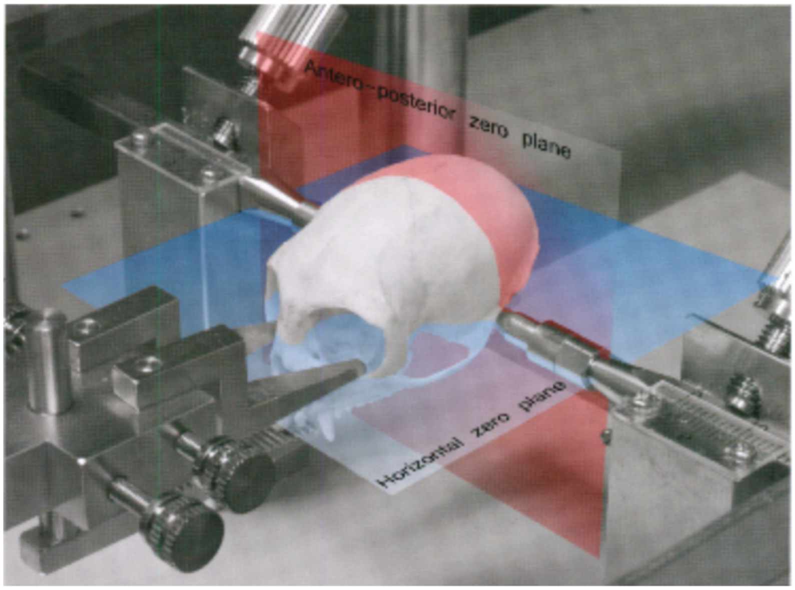


##### Coordinate directions

We use an RAS (right, anterior, superior) coordinate system where the "x" direction points from left to right, the "y" direction points from posterior to anterior, and the "z" direction points from inferior to superior.

##### Coordinate origin

In the common marmoset, the horizontal zero plane is defined as the plane passing through the lower margin of the orbit and the center of the external auditory meatus (see figure) (note you cannot see bregma on marmoset, it is fused too tightly to see). In an imaging apparatus, the skull is fixed through the ears, so this is a good choice. The anteroposterior zero plane is defined as the plane perpendicular to the horizontal zero plane which passes the centers of the external auditory meati. The left-right zero plane is the midsagittal plane (Saavedra and Mazzuchelli, 1969; Stephan et al., 1980).

###### Other information

**Note:** On our CSH dropbox system the atlas data is located here: https://www.dropbox.com/sh/70hg40e8b3ro9vx/AACk05Hm-BFGbMD5NIQolqtxa?dl=0 with images:

Nissl reference: bma-1-nissl.nii.gz

MRI reference: bma-1-mri.nii.gz

Atlas: bma-1-region_seg.nii.gz

**Note:** We have built a population average image for males and females, which is located on CSH at /nfs/data/main/M38/marmoset_ccf

##### 3.1.3 Human atlas

We use the Montreal Neurological Institute - International Consortium for Brain Mapping (MNI-ICBM) coordinate system, which is described here: https://www.mcgill.ca/bic/software/tools-data-analysis/anatomical-mri/atlases/icbm152-non-linear . An example is shown below.

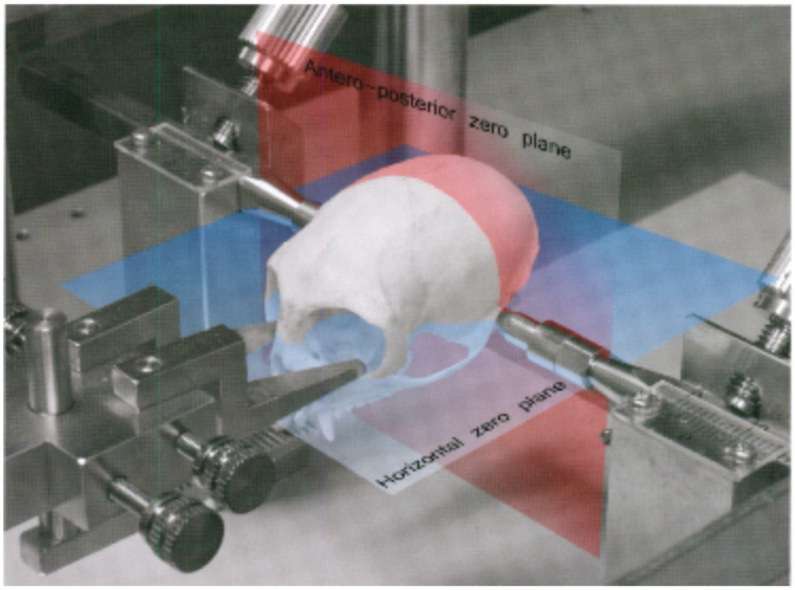


###### Coordinate directions

The atlas is based on the Talarach Tournoux coordinate system which is RAS. "x" points from left to right, "y" points from posterior to anterior (posterior commissure to anterior commissure), and "z" points from inferior to superior.

###### Coordinate origin

The coordinate origin is 4 millimeters above the center of the anterior commissure.

Details: these points are intersection of planes. AC to PC defines a line. Left right defines a line. These two lines define a plane.

Midsagittal plane defines another plane.

The intersection defines a line, and the left right center on this line gives the origin.

###### Voxel size

Standard voxel size is 1mm isotropic, but other possibilities are available.

###### Other information

Currently can be downloaded form the web at the above link https://www.mcgill.ca/bic/software/tools-data-analysis/anatomical-mri/atlases/icbm152-non-linear .

**Note:** Another possibility is MNI-Colin27 and not ICBM152 nonlinear. https://www.mcgill.ca/bic/software/tools-data-analysis/anatomical-mri/atlases/colin-27-2008

Note that the difference is the 4mm offset.

#### 3.2 Input space

The input space describes the coordinate system of 2D serial section images to be registered. This is the internal structure of the pipeline, but preprocessors may be used to convert data with other structures. For example, point sets may be described by the integer index of a pixel in an image, rather than a location with units of length. A converter is necessary in this case (for example `emlddmm.convert_points_from_json()`)

##### 3.2.1 Coordinate directions

The "x" coordinate points from left to right (i.e. from the left side to the right side of a 2D image displayed on a screen). The "y" coordinate points from up to down. The "z" coordinate points from the first acquired slice to the last acquired slice.

**Note:** This convention does not reference any anatomy, only the camera. This is chosen because sections could have any orientation (coronal, sagittal, etc.).

##### 3.2.2 Coordinate origin

The xy origin will always be in the center of the image. i.e. On a given slice if you find the average "x" coordinate, or the average "y" coordinate, it will be 0. The z coordinate origin is also chosen such that slices are centered for this dataset: the average of the z coordinate for the first and last slices is zero.

**Note:** Motivation for this choice is that we can pad images symmetrically without changing the origin, and rotation about this origin is more numerically stable than rotation around one corner.

##### 3.2.3 Voxel size

Voxel size is an input parameter to the pipeline (e.g. stored in json sidecar files for 2D images, or in vtk headers for 3D images). Our convention is to use units of microns.

**Note:** In a typical workflow, an image is created with resolution 0.46umx0.46micron, with a slice thickness of 10 micron. For registration purposes these are typically downsampled by a factor of 32 in the x and y directions, making the resolution 14.72um before they are input to the pipeline.

#### 3.3 Registered space

In our workflow, any time a 3D to sequence of 2D slices map is calculated, a new registered space is created (see for example *here*). A sequence of rigid transformations are applied to each 2D slice to match the general shape of a deformed reference 3D image. This effectively defines a 3D coordinate space.

##### 3.3.1 Coordinate directions

Same as *input space directions*.

##### 3.3.2 Coordinate origin

Same as *input space origin*.

##### 3.3.3 Voxel size

Same as *input space voxel size*.

**Note:** Once transformations have been computed, high resolution data is transformed into registered space for display on the web or other viewers. This high resolution data uses its native voxel size (typically 0.46 microns).

##### 3.3.4 The non-uniqueness of registered space

Our mapping algorithm enforces alignment between data in the common space, and data in the input space. This sequence of transforms can be factored to define a space in the "middle". This is a factorization of transformations problem. Just like matrix factorizations are not unique without constraints, the data does not uniquely define a registered space. Rather, we use several heuristics to define a space that is a "minimally distorted" version of the common space. The general idea is that any component of the transform that can be represented as a sequence of 2D transforms, should be represented that way, rather than as part of a 3D transform. In particular

- We want no translation in the xy direction of the 3D affine transformation. (this is enforced by our pipeline)
- We want no shear perpendicular to the z axis in the 3D transformation. (this can be enforced by chosing to limit the affine transformation to rigid, or rigid plus scale)
- After applying our affine transform we want the up vector to still point up, when projected into the slice plane. (implemented with the `up_vector` parameter in `emlddmm.emlddmm()`)

##### 3.3.5 Other information

While registered space uses the same coordinate system conventions as input space, the set of voxel locations images should be sampled on is likely to be different. One example occurs when our atlas has its origin at the bregma point on the skull, but our input data has its origin in the center of the brain tissue. Due to our factorization conventions described *above*, tissue in registered space will no longer be centered at the origin.

In a typical situation, our input histology is sectioned in either the coronal, sagittal, or transverse plane, and we map it to a well characterized atlas. In these situations, the origin in xy for the registered space can be interpretted with respect to the atlas origin, and will correspond to the origin for two of the 3 axes in the atlas. We enumerate sevaral cases below and provide some information explicitly.

###### Mouse with coronal sections

Using our *Mouse atlas*, and a coronally sectioned dataset in *Input space*, input space x corresponds to the right left axis, and input space y corresponds to the dorsal ventral axis.

Therefore, the x=0 point in registered space corresponds to the anatomy at the y=0 point in the atlas, and the y=0 point in registered space corresponds to the anatomy at the x=0 point in the atlas.

When reconstructing imaging data in this space we chose a set of sample points for voxels that will cover the anatomy. Therefore, we sample x starting at -5695.0 um, ending at 5695.06, and using 24762 samples equally spaced by 0.46 microns. Similarly, we sample y starting at -870.0 um, ending at 7120.2 um, and using 17371 samples equally spaced by 0.46 microns.

This convention allows us to convert between spatial locations and pixel indices (row=i,col=j, starting at 0), using the follow formulas:

$$i = \text{round}[(y - (-870.0))/0.46]$$
$$j = \text{round}[(x - (-5695.0))/0.46]$$
$$y = 0.46i + (-870.0)$$
$$x = 0.46j + (-5695.0)$$

##### Mouse with sagittal sections

Using our *Mouse atlas*, and a sagittaly sectioned dataset in *Input space*, input space x corresponds to the anterior posterior axis (note the nose will be on the left), and input space y corresponds to the dorsal ventral axis.

Therefore, the x=0 point in registered space corresponds to the anatomy at the z=0 point in the atlas, and the y=0 point in registered space corresponds to the anatomy at the x=0 point in the atlas.

When reconstructing imaging data in this space we chose a set of sample points for voxels that will cover the anatomy. Therefore, we sample x starting at -7970.0 um, ending at 5220.04, and using 28675 samples equally spaced by 0.46 microns. Similarly, we sample y starting at -870.0 um, ending at 7120.2 um, and using 17371 samples equally spaced by 0.46 microns.

This convention allows us to convert between spatial locations and pixel indices (row=i,col=j, starting at 0), using the follow formulas:

$$
\begin{aligned}
i &= \text{round}[(y - (-870.0))/0.46] \\
j &= \text{round}[(x - (-7970.0))/0.46] \\
y &= 0.46i + (-870.0) \\
x &= 0.46j + (-7970.0)
\end{aligned}
$$

##### Mouse with transverse sections

Using our *Mouse atlas*, and a transverse sectioned dataset in *Input space*, input space x corresponds to the anterior posterior axis (CHECK!), and input space y corresponds to the right left axis.

Therefore, the x=0 point in registered space corresponds to the anatomy at the z=0 point in the atlas, and the y=0 point in registered space corresponds to the anatomy at the y=0 point in the atlas.

When reconstructing imaging data in this space we chose a set of sample points for voxels that will cover the anatomy. Therefore, we sample x starting at -7970.0 um, ending at 5220.04, and using 28675 samples equally spaced by 0.46 microns. Similarly, we sample y starting at -5695.0 um, ending at 5695.09 um, and using 24762 samples equally spaced by 0.46 microns.

This convention allows us to convert between spatial locations and pixel indices (row=i,col=j, starting at 0), using the follow formulas:

$$
\begin{aligned}
i &= \text{round}[(y - (-5695.0))/0.46] \\
j &= \text{round}[(x - (-7970.0))/0.46] \\
y &= 0.46i + (-5695.0) \\
x &= 0.46j + (-7970.0)
\end{aligned}
$$

CHAPTER

FOUR

### FILE FORMATS

We propose to use VTK formatted data whenever possible. Currently we use simple legacy .vtk files (https://docs.vtk.org/en/latest/design_documents/VTKFileFormats.html). This supports vector and raster graphics, works well with visualization software including web viewers, and is largely human readable. It does not support compression, and so other formats are also used.

#### 4.1 JSON geometry files

Every imaging file that does not store geometry information (e.g. 2D slices stored as pngs/tifs/etc) will have a corresponding short JSON sidecar file located in the same directory, with the same name filename, and the extension .json appended. The information stored here should be as close as possible to an NRRD header (http://teem.sourceforge.net/nrrd/format.html). Note that 2D images are described as though they are 3D, with pixel size in the z dimension referring to section thickness.

Such sidecar files are inspired by the BIDS standard, and contain information typically stored in an NRRD header. Each sidecar file must contain the following fields:

- "DataFile": the image file name
- "SpaceDirections": a list of vectors for each image dimension specifying the unit conversion from pixel indices (row,column) to input space coordinates. Note that the z-coordinate conversion indicates the slice thickness plus the spacing between slices. This is a list of vectors in x,y,z order, where the xyz coordinate system is defined in the "input space" section.
- "SpaceOrigin": The world coordinate of the image origin, in x,y,z order.

Other metadata can be stored in the json file, but only the above 3 are generally used for the data loader functions.

Note that each 2D image is modeled as a 3D image with a single slice (i.e. a size of 1 in).

An example is shown below:

```
{
  "DataFile": "MD787_small_nissl/MD787-N27-2019.03.28-22.55.54_MD787_2_0080.png",
  "Type": "Float32",
  "Dimension": 3,
  "Endian": "big",
  "Sizes": [3, 392, 480, 1],
  "Space": "inferior-right-posterior",
  "SpaceDimension": 3,
  "SpaceUnits": ["um", "um","um" ],
  "SpaceDirections": [
    "none",
```

(continues on next page)

(continued from previous page)

```
    [44.160000000000004, 0.0, 0.0],
        [0.0, 44.160000000000004, 0.0],
        [0.0, 0.0, 200      ]
  ],
  "SliceThickness" : 10.0
  "SpaceOrigin": [-8633.28, -10576.32, -120100.0]
}
```

Note when data is read, the space directions are reversed to line up with common conventions for image array axes. The last axis of an image array corresponds to columns (x), the second last corresponds to rows (y), and the third last corresponds to slices (z).

#### 4.2 Dataset lists

Since sections may be missing or require other comments, we include a tsv file in the same directory describing every slice in the dataset. The required fields are sample_id and status, the latter should contain present or absent. An example is shown below:

```
sample_id    participant_id          species         status
MD787-N7-2019.03.28-22.05.43_MD787_2_0020.png       MD787   Mus Musculus    present
MD787-N14-2019.03.28-22.20.46_MD787_1_0040.png      MD787   Mus Musculus    present
MD787-N20-2019.03.28-22.36.39_MD787_3_0060.png      MD787   Mus Musculus    present
MD787-N27-2019.03.28-22.55.54_MD787_2_0080.png      MD787   Mus Musculus    present
MD787-N34-2019.03.28-23.15.58_MD787_1_0100.png      MD787   Mus Musculus    present
MD787-N40-2019.03.28-23.33.43_MD787_3_0120.png      MD787   Mus Musculus    present
MD787-N47-2019.03.28-23.54.40_MD787_2_0140.png      MD787   Mus Musculus    present
MD787-N54-2019.03.29-00.15.46_MD787_1_0160.png      MD787   Mus Musculus    present
MD787-N60-2019.03.29-00.33.42_MD787_3_0180.png      MD787   Mus Musculus    present
MD787-N67-2019.03.29-00.56.05_MD787_2_0200.png      MD787   Mus Musculus    present
MD787-N74-2019.03.29-01.18.34_MD787_1_0220.png      MD787   Mus Musculus    present
MD787-N80-2019.03.29-01.36.50_MD787_3_0240.png      MD787   Mus Musculus    present
MD787-N87-2019.03.29-01.57.37_MD787_2_0260.png      MD787   Mus Musculus    present
MD787-N94-2019.03.29-02.19.41_MD787_1_0280.png      MD787   Mus Musculus    present
MD787-N100-2019.03.29-02.40.34_MD787_3_0300.png     MD787   Mus Musculus    present
MD787-N107-2019.03.29-03.04.17_MD787_2_0320.png     MD787   Mus Musculus    present
MD787-N114-2019.03.29-03.28.07_MD787_1_0340.png     MD787   Mus Musculus    present
MD787-N120-2019.03.29-03.49.00_MD787_3_0360.png     MD787   Mus Musculus    present
```

#### 4.3 Legacy CSV geometry files

In older versions of our pipeline, we stored information in a csv file, with the following 10 fields for each image.

- Filename
- Nx ny nz: number of pixels in the x y and z direction (nz=1 if 2D image)
- Dx dy dz: pixel size in x y and z direction (dz = slice thickness if 2D image)
- X0 y0 z0: coordinate of the first pixel in x y and z direction (z0 = location within dataset if 2D image)

#### 4.4 Cold Spring Harbor legacy geometry files

Cold Spring Harbor is storing geometry data in a plain text file. For example:

```
2021-10-05 14:30:13.177668
Registered : Y
Input Path:/nfs/data/main/M32/RegistrationData/Data_Marmoset/m6344/Transformation_OUTPUT/
↪m6344_img/
Output Path:/nfs/data/main/M32/Cell_Detection/CellDetPass1_reg/m6344/
Number of Files Detected:386
Resolution:0.92
Resolution in Json: 1 micron/pixel
```

#### 4.5 3D imaging data

Our standard is to use simple vtk legacy format for 3D (see https://examples.vtk.org/site/VTKFileFormats/#simple-legacy-formats). Note that this data is always stored in big endian, regardless of machine defaults. These have simple human readable headers that contain the 9 pieces of information above. Our pipeline provides basic support for nifti images using the `nibabel` python package.

#### 4.6 2D microscopy images from Cold Spring Harbor

Acquired microscopy data is stored at Cold Spring Harbor Laboratory in jp2 format at full resolution (generally 0.46 microns per pixel). The filename is generated by the scanner, following a template schema that the Mitra lab uses in a standard manner. An example is:

```
MD787-N3-2019.03.28-21.57.34_MD787_3_0009.jp2
```

Where the meaning of each hyphen separated field is:

```
{sample id}-{N/F/IHC for nissl fluoro or ihc}-{slide number}-{date}-{time}_{sample id}_
↪{what position on slide}_{section number id in anterior to posterior order (generally)}
↪.
```

Note that no geometry information is stored in filenames here, so this should be added as a json companion file.

#### 4.7 2D datasets for registration

Typically data is downsampled by a factor of 32 and saved as a .tif with the same filename.

Registration data can be safely downsampled to approximately the same resolution as atlas images (10-50 micron).

For 2D serial section datasets images should be stored in a single directory using standard imaging formats (i.e. to be read by matplotlib's imread function), downsampled by approximately 32 times (e.g. 14.72 microns for CSH data). While our pipelines do support downsampling to desired resolutions, code will run more efficiently if these sections are already downsampled.

Slice datasets must contain sidecar json files, and data set list tsv files. The script, `histsetup`, generates sidecar files and dataset lists given a subject dataset and voxel spacing (where spacing in the z axis indicates slice thickness plus slice spacing).

#### 4.8 Affine Transformations

Affine transformations are stored as 4x4 matrices written in a text file. Each column is separated by spaces. Each row is separated by a new line. Coordinates are in xyz order. When read into python using our library, they will be converted to zyx order to be consistent with our conventions for indexing image arrays.

#### 4.9 Deformations

Deformations are as 3 component displacement fields (not position fields) in vtk files. In python we work in zyx order, but when writing to vtk fields we switch to xyz order which is the vtk convention.

#### 4.10 Velocity fields

Velocities are nt x 3 component vector fields in vtk files. In python we work in zyx order, but when writing to vtk fields we switch to xyz order which is the vtk convention.

#### 4.11 Annotations

2D annotations are stored as geojson files using the multipolygon data type. Each structure is given a name, and an integer ID. Metadata stores information about which atlas is used, and which 2D image file the annotations correspond to.

These files will also contain atlas coordinate gridlines.

#### 4.12 Point sets

Point sets are stored in vtk polydata format.

#### 4.13 Pixel indexed point sets

Point sets that describe detected cells are stored in geojson format. These point sets have some constraints based on how they will be displayed using open layers or angular on the web.

CHAPTER

FIVE

### INPUT SPECIFICATION VIA TRANSFORMATION GRAPH INTERFACE

We support pipelines for registering several datasets to each other, and reconstructing data from one dataset in the space of any other dataset. All of the registrations and reconstructions can be performed by executing a single command from the command line with one input, a json file which contains the following information:

#### 5.1 Names of spaces

Registrations are computed between pairs of spaces. Each space should be given a unique name. (e.g. “atlas”, “CT”, “exvivoMRI”,”invivoMRI”, “Histology”).

#### 5.2 Names of images

Each space may have more than one imaging dataset sampled in it (for example multiple MRI scans with different contrasts). Each image within a space should be given a unique name. (e.g. “exvivoMRI -> T1”, “exvivoMRI -> T2”, “invivoMRI -> T1”, “Histology”)

#### 5.3 Filenames

Each image should have a filename (for 3D data), or a directory (for 2D data) associated with it.

#### 5.4 Registration tuples

To register a complex multimodal dataset, we specify a list of (space/image to map from, space/image to map to ) tuples. These correspond to edges in a graph and should span the set of spaces. This set of transformations will be computed using our optimization procedure.

#### 5.5 Registration Configurations

Each registration is computed using unique parameters specified in a registration configuration json file whose path must be listed. These will be loaded into python into a dictionary, which will be passed to functions via keyword arguments. Each registration is run in a multi scale fashion, from low resolution to high resolution, and so each parameter should be specified as a list (one entry for each resolution) or a singleton list (one entry for all resolutions). We have included examples of registration config files in the examples folder.

#### 5.6 Reconstruction tuples

After transformations are computed, we can reconstruct data from one space in any other space. Tuples of the form (space/image to map from, space to map to) are specified. Given the registration tuples, a path of transformations will be computed, which may involve the composition of more than one calculated transform. We can also choose to reconstruct each image in every other space instead of specifying each mapping with a tuple.

#### 5.7 Example

For example we can run registration and reconstruction with the command:

```
python transformation_graph.py --infile INPUT_JSON_FILE
```

Where the input json file contains:

```
{
    "space_image_path": [["MRI", "masked", "/home/brysongray/data/MD816_mini/HR_NIHxCSHL_
↪50um_14T_M1_masked.vtk"],
                        ["CCF", "average_template_50", "/home/brysongray/data/MD816_mini/
↪average_template_50.vtk"],
                        ["MRI", "unmasked", "/home/brysongray/data/MD816_mini/HR_NIHxCSHL_
↪50um_14T_M1.vtk"],
                        ["CT", "masked", "/home/brysongray/data/MD816_mini/ct_mask.vtk"],
                        ["HIST", "nissl", "/home/brysongray/data/MD816_mini/MD816_STIF_
↪mini"]],
    "registrations": [[["MRI", "masked"], ["HIST", "nissl"] ],
                        [ ["CCF", "average_template_50"], ["MRI", "masked"],],
                        [["CT", "masked"], ["MRI", "masked"] ]],
    "configs": ["/home/brysongray/emlddmm/config787small.json",
                "/home/brysongray/emlddmm/configMD816_MR_to_CCF.json",
                "/home/brysongray/emlddmm/configMD816_MR_to_CT.json"],
    "output": "/home/brysongray/emlddmm/transformation_graph_outputs",
    "transforms": [[["CCF", "average_template_50"], ["HIST", "nissl"] ],
                  [["CT", "masked"], ["MRI", "masked"] ]],
    "transform_all": "False"
}
```

This input structure will do the following:

1. It will define 4 spaces, called MRI, CCF, CT and HIST
2. It will define images in these spaces. Paths to images are provided.
    - Two images in the MRI space, called “masked” and “unmasked”.

    - It will define one image in CCF space called "average_template_50".
    - It will define one image in CT space called "masked".
    - It will define one image set in HIST space, called "nissl".

3. It will define a set of registrations to calculate. Each registration requires a pair of spaces, and an image name within that space.
    - It will registered the masked MRI to the histology.
    - It will register the CCF atlas to the masked MRI
    - It will register the masked CT to the masked MRI
4. Each registration will be calculated in order, using the config files provided for parameters.
5. Outputs of the registration processes will be saved in the specified output directory.
6. Calculated transforms are applied to map images into new spaces
    - The average template is mapped into the HIST space
    - The masked CT images is mapped into the MRI space.
7. Generally we reconstrct all images in all spaces, in which case transform_all is set to true.

The registration procedure internally sets up the following graph

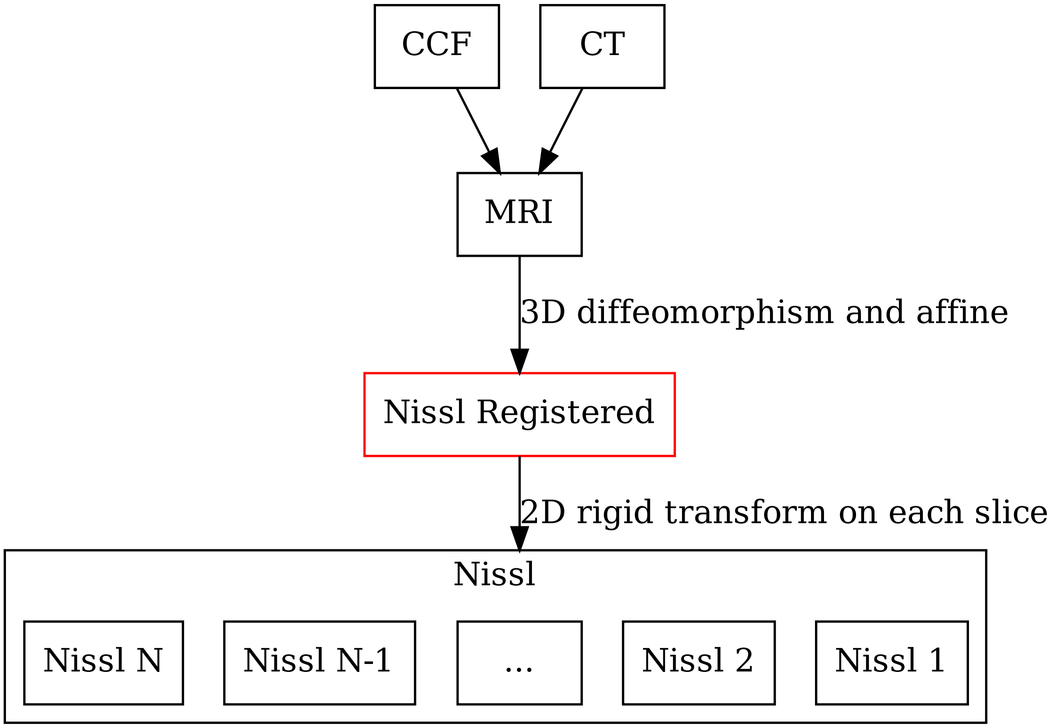


Fig. 1: For any 3D to 2D map, a registered space is automatically created (shown in red). No input data is associated with this space, but images can be reconstructed into this space.

CHAPTER

SIX

### OUTPUT SPECIFICATION

Our output data structure contains transformations between pairs of named spaces (always), transformed images (suggested but not necessary), and other data types such as points and geojson annotations.

These pairs are organized in a hierarchical tree, where the parent directories contain data in a given space, and the child directories contain data from a given space.

#### 6.1 Example

Example output data structure is shown here. Lists are used to show directory hierarchy: This supports an arbitrary number of folders.:

```
{Space i}
  {Space j}_to_{space i}
    Transforms (always)
      {space i}_to_{space j}_displacement.vtk (3D to 3D, or 3D to registered space, NOT␣
↪3D to input which does not exist as a displacement field)
      {space i}_{image k}_to_{space j}_{image k'}_matrix.txt (2D to 2D only)
      {space i}_{image k}_to_{space j}_displacement.vtk (i 2D to 3D only)
    Images (suggested)
      {space j}_{image k}_to_{space i}.vtk
      {space j}_{image k}_to_{space i}_{image k'}.vtk (for 2D to 2D)
    Points (optional)
      {space j}_{image k}_detects_to_{space i}.vtk
    Json (for atlas only)
      Atlas_to_{space j}_{image k}.geojson
    Meanxyz (for atlas only)
      {space j}_{image k}_detects_to_atlas_meanxyz.txt
  QC (optional)
    Composite_{image slice name}_QC.jpg
```

#### 6.2 Notes

Some important notes are below:

1. Output raster data is stored using simple legacy vtk file format (see *here*).
2. Output point data is stored using simple legacy vtk file format, with polydata.
3. json is shown only for data from atlas to a 2D space.
4. Mean xyz is shown only for a 2D space to the atlas.
5. Transforms are stored as a rigid transformation matrix only for maps from a 2D space to another 2D space.
6. Note the "to" in the naming of transforms is opposite to images. This is intentional.
7. Note that in 2D directories, image names are appended to space names for uniqueness, separated by an underscore.
8. QC figures are not standard, as they will vary by dataset.

CHAPTER

SEVEN

### EXAMPLES

We include two worked examples in our github repository under the examples folder. In both cases we include an interactive approach that inclues several visualizations, in the form of a jupyter notebook. At the end of each jupyter notebook we show how the same analysis can be run from the command line, using our transformation graph interface, producing our standard output format.

The first describes 3D registration between two human MRI datasets. The second describes 3D to 2D registration between the Allen Nissl atlas, and a sequence of Nissl stained images.

Note that our command line interface has only been validated on Linux systems.

#### 7.1 Human MRI example

In this example we register a pair of 3D human brain MR images.

First we walk through the example in this notebook.

Then we write config files to disk, and run the example from the command line. The command line interface has only been validated on Linux systems.

##### 7.1.1 Import libraries

[1]:
```
# numpy for multidimensional arrays to store images
import numpy as np
# matplotlib for visualization
import matplotlib.pyplot as plt
# the command below will allow interactive figures that update as code runs
%matplotlib notebook

# import tools for working with files
from os import makedirs
from os.path import join

# import the json library for writing out config files
import json

# import the subprocess library for running code from command line
import subprocess

# import the emlddmm image registration library
```

(continues on next page)

(continued from previous page)

```
import sys
sys.path.append('../..')
import emlddmm
```

##### 7.1.2 Outputs

[2]:
```
output_directory = 'human_mri_example_notebook_outputs'
```

[3]:
```
makedirs(output_directory,exist_ok=True)
```

##### 7.1.3 Load images

[4]:
```
target_name = 'TargetMRI.vtk'
atlas_name = 'AtlasMRI.vtk'
label_name = 'AtlasLabels.vtk'
```

[5]:
```
# load the atlas with normalization (mean of abs is 1)
xI,I,_,_ = emlddmm.read_data(atlas_name,normalize=True)
# draw a picture
fig,ax = emlddmm.draw(I,xI,cmap='gray')
fig.suptitle('Atlas image')
```

```
<IPython.core.display.Javascript object>
```

```
<IPython.core.display.HTML object>
```

[5]:
```
Text(0.5, 0.98, 'Atlas image')
```

[6]:
```
# load the atlas segmentation labels, with no normalization (because these are integer␣
↪labels)
xS,S,_,_ = emlddmm.read_data(label_name)
SRGB = emlddmm.labels_to_rgb(S,black_label=256)
# draw a picture, showing labels and MRI
fig,ax = emlddmm.draw(SRGB+I/np.max(I)*2.0,xS)
fig.suptitle('Atlas image')
```

```
<IPython.core.display.Javascript object>
```

```
<IPython.core.display.HTML object>
```

[6]:
```
Text(0.5, 0.98, 'Atlas image')
```

[7]:
```
# load the target with normalization (mean of abs is 1)
xJ,J,_,_ = emlddmm.read_data(target_name,normalize=True)
# draw a picture
fig,ax = emlddmm.draw(J,xJ,cmap='gray')
fig.suptitle('Target image image')
```

```
<IPython.core.display.Javascript object>
```

```
<IPython.core.display.HTML object>
```

[7]:
```
Text(0.5, 0.98, 'Target image image')
```

[8]:
```
# run registration at 3 different spatial scales
```

[9]:
```
config = {
    'device':'cpu', # cpu or cuda:0
    'downI':[[4,4,4],[2,2,2],[1,1,1]], # downsampling factors for atlas at multiple␣
↪scales
    'downJ':[[4,4,4],[2,2,2],[1,1,1]], # downsampling factors for target at multiple␣
↪scales
    'n_iter':[50,40,30], # how many iterations of gradient descent at each scale
    'v_start':[0], # at what iteration of gradient descent do we start optimizing over␣
↪deformation
    'eA': [1e1], # gradient descent stepsize for 3D affine transform
    'ev':[5e-1], # gradient descent stepsize for the deformation
    'a':2.0, # spatial scale of deformation
    'dv':2.0, # sampling interval for deformation
    'sigmaR':5e0, # regularizatoin for deformation (bigger means less regularization)
    'local_contrast':[[32,32,32]] # divide the images into small blocks to estimate␣
↪contrast differences
}
```

[10]:
```
out = emlddmm.emlddmm_multiscale(xI=[xI],I=I,xJ=[xJ],J=J,**config)
```

```
Found 3 scales
```

```
<IPython.core.display.Javascript object>
```

```
<IPython.core.display.HTML object>
```

```
<IPython.core.display.Javascript object>
```

```
<IPython.core.display.HTML object>
```

```
<IPython.core.display.Javascript object>
```

```
<IPython.core.display.HTML object>
```

```
<IPython.core.display.Javascript object>
```

```
<IPython.core.display.HTML object>
```

```
<IPython.core.display.Javascript object>
```

```
<IPython.core.display.HTML object>
```

```
<IPython.core.display.Javascript object>
```

```
<IPython.core.display.HTML object>
```

```
<IPython.core.display.Javascript object>
```

```
<IPython.core.display.HTML object>
```

```
<IPython.core.display.Javascript object>
```

```
<IPython.core.display.HTML object>
```

```
../../emlddmm.py:175: RuntimeWarning: invalid value encountered in true_divide
  J /= (vmax[:,None,None,None] - vmin[:,None,None,None])
```

```
Iteration 30, linear oscilating, reducing eA to 9.0
Iteration 40, translation oscilating, reducing eA to 8.1
```

```
../../emlddmm.py:1334: UserWarning: To copy construct from a tensor, it is recommended␣
↪to use sourceTensor.clone().detach() or sourceTensor.clone().detach().requires_grad_
↪(True), rather than torch.tensor(sourceTensor).
  v = torch.tensor(v.detach().clone(),device=device,dtype=dtype)
../../emlddmm.py:1378: UserWarning: To copy construct from a tensor, it is recommended␣
↪to use sourceTensor.clone().detach() or sourceTensor.clone().detach().requires_grad_
↪(True), rather than torch.tensor(sourceTensor).
  A = torch.tensor(A.detach().clone(),device=device,dtype=dtype)
```

```
<IPython.core.display.Javascript object>
```

```
<IPython.core.display.HTML object>
```

```
<IPython.core.display.Javascript object>
```

```
<IPython.core.display.HTML object>
```

```
<IPython.core.display.Javascript object>
```

```
<IPython.core.display.HTML object>
```

```
<IPython.core.display.Javascript object>
```

```
<IPython.core.display.HTML object>
```

```
<IPython.core.display.Javascript object>
```

```
<IPython.core.display.HTML object>
```

```
<IPython.core.display.Javascript object>
```

```
<IPython.core.display.HTML object>
```

```
<IPython.core.display.Javascript object>
```

```
<IPython.core.display.HTML object>
```

```
<IPython.core.display.Javascript object>
```

```
<IPython.core.display.HTML object>
```

```
<IPython.core.display.Javascript object>
```

```
<IPython.core.display.HTML object>
```

```
../../emlddmm.py:1445: RuntimeWarning: More than 20 figures have been opened. Figures␣
↪created through the pyplot interface (`matplotlib.pyplot.figure`) are retained until␣
↪explicitly closed and may consume too much memory. (To control this warning, see the␣
↪rcParam `figure.max_open_warning`).
  figA,axA = plt.subplots(2,2)
```

```
<IPython.core.display.Javascript object>
```

```
<IPython.core.display.HTML object>
```

```
<IPython.core.display.Javascript object>
```

```
<IPython.core.display.HTML object>
```

```
<IPython.core.display.Javascript object>
```

```
<IPython.core.display.HTML object>
```

```
<IPython.core.display.Javascript object>
```

```
<IPython.core.display.HTML object>
```

```
<IPython.core.display.Javascript object>
```

```
<IPython.core.display.HTML object>
```

```
<IPython.core.display.Javascript object>
```

```
<IPython.core.display.HTML object>
```

```
<IPython.core.display.Javascript object>
```

```
<IPython.core.display.HTML object>
```

##### 7.1.4 apply the transform to the atlas image

We use the backward (inverse) transformations.

[11]:
```
tform = emlddmm.compose_sequence(
    [
        emlddmm.Transform(out[-1]['A'],direction='b'),
        emlddmm.Transform(out[-1]['v'],domain=out[-1]['xv'],direction='b')
    ],
    xJ
)
AphiI = emlddmm.apply_transform_float(xI,I,tform)
AphiS = emlddmm.apply_transform_int(xS,S,tform)
```

[12]:
```
# draw the labels over the target image
AphiSRGB = emlddmm.labels_to_rgb(AphiS,black_label=256)
# draw a picture, showing labels and MRI
fig,ax = emlddmm.draw(AphiSRGB+J/np.max(J)*2.0,xJ)
fig.suptitle('Target image with atlas labels')
```

```
<IPython.core.display.Javascript object>
```

```
<IPython.core.display.HTML object>
```

[12]:
```
Text(0.5, 0.98, 'Target image with atlas labels')
```

[13]:
```
# save the transformed images
emlddmm.write_data(join(output_directory,'atlas_labels_to_target.vtk'),xJ,AphiS,title=
↪'atlas_labels_to_target')
emlddmm.write_data(join(output_directory,'atlas_image_to_target.vtk'),xJ,AphiI,title=
↪'atlas_image_to_target')
```

##### 7.1.5 apply the transform to the target image

We use the forward transformation

[14]:
```
# transform target back to template
tform = emlddmm.compose_sequence(
    [
        emlddmm.Transform(out[-1]['v'],domain=out[-1]['xv'],direction='f'),
        emlddmm.Transform(out[-1]['A'],direction='f'),
    ],
    xI
)
phiiAiJ = emlddmm.apply_transform_float(xJ,J,tform)

# resample target on template voxels
tform = emlddmm.compose_sequence([emlddmm.Transform(np.eye(4))],xI)
Jresampled = emlddmm.apply_transform_float(xJ,J,tform)
```

[15]:
```
fig,ax = emlddmm.draw(np.concatenate((I,Jresampled)),xI)
fig.suptitle('Atlas in magenta, UNtransformed target in green')

fig,ax = emlddmm.draw(np.concatenate((I,phiiAiJ)),xI)
fig.suptitle('Atlas in magenta, transformed target in green')
```

```
<IPython.core.display.Javascript object>
```

```
<IPython.core.display.HTML object>
```

```
<IPython.core.display.Javascript object>
```

```
<IPython.core.display.HTML object>
```

[15]:
```
Text(0.5, 0.98, 'Atlas in magenta, transformed target in green')
```

[16]:
```
# save the transformed image
emlddmm.write_data(join(output_directory,'target_image_to_atlas.vtk'),xJ,AphiS,title=
↪'target_image_to_atlas')
```

##### 7.1.6 Run the same analysis using the command line interface

Next we will show how to produce necessary config files, and run the same example using our transformation graph command line interface.

##### 7.1.7 write out a config file for registration

[17]:
```
config_file = 'atlas_to_target_config.json'
with open(config_file,'wt') as f:
    json.dump(config,f)
```

##### 7.1.8 write out a config file for the transformation graph

[18]:
```
transformation_config = {
  "output": "output",
  "space_image_path": [
    [
      "Atlas",
      "image",
      atlas_name
    ],
    [
      "Atlas",
      "labels",
      label_name
    ],
    [
      "Target",
      "image",
      target_name
    ],
  ],
  "registrations": [
    [
      [
        "Atlas",
        "image"
      ],
      [
        "Target",
        "image"
      ]
    ],
  ],
  "configs": [
    config_file,

  ],
  "transform_all":True,
}
transformation_config_file = 'transformation_graph_config.json'
with open(transformation_config_file,'wt') as f:
    json.dump(transformation_config,f)
```

##### 7.1.9 Run this example from the command line

We will write out the parameters above to a config file, and run our command line interface. This will produce all our standard outputs.

```
[19]: command = f'python -u ../../transformation_graph_v01.py --infile {transformation_config_
      ↪file} > outputs.txt 2>&1'
      print('about to run command:')
      print(command)
```

```
about to run command:
python -u ../../transformation_graph_v01.py --infile transformation_graph_config.json >␣
↪outputs.txt 2>&1
```

```
[20]: subprocess.call(command,shell=True)
```

```
[20]: 0
```

##### 7.1.10 View all the outputs

All the output directories are printed below. Note that this includes a python file graph.p, which contains information about the transformation graph. This will allow us to apply the transforms we have already calculated to new datasets later.

```
[27]: from os import walk
      for dirpath,dirnames,filenames in walk('output',):
          for f in filenames:
              print(join(dirpath,f))
```

```
output/infile.json
output/graph.p
output/Atlas/Target_to_Atlas/transforms/velocity.vtk
output/Atlas/Target_to_Atlas/transforms/A.txt
output/Atlas/Target_to_Atlas/transforms/Target_to_Atlas_displacement.vtk
output/Atlas/Target_to_Atlas/transforms/Target_image_to_Atlas_detjac.vtk
output/Atlas/Target_to_Atlas/qc/Target_image_to_Atlas.jpg
output/Atlas/Target_to_Atlas/qc/Atlas_image.jpg
output/Atlas/Target_to_Atlas/images/Target_image_to_Atlas.vtk
output/Target/Atlas_to_Target/qc/Atlas_image_to_Target.jpg
output/Target/Atlas_to_Target/qc/Target_image.jpg
output/Target/Atlas_to_Target/images/Atlas_image_to_Target.vtk
output/Target/Atlas_to_Target/images/Atlas_labels_to_Target.vtk
output/Target/Atlas_to_Target/transforms/Atlas_to_Target_displacement.vtk
output/Target/Atlas_to_Target/transforms/Atlas_image_to_Target_detjac.vtk
output/Target/Atlas_to_Target/transforms/Atlas_labels_to_Target_detjac.vtk
```

#### 7.2 Mouse Serial Section Example

In this example we will register the Allen CCF atlas to a mouse Nissl dataset.

First we will walk through an exapmle in this notebook.

Then we write config files to disk, and run the example from the command line. The command line interface has only been validated on Linux systems.

##### 7.2.1 Import libraries

[1]:
```
# numpy for multidimensional arrays to store images
import numpy as np
# matplotlib for visualization
import matplotlib.pyplot as plt
# the command below will allow interactive figures that update as code runs
%matplotlib notebook

# import tools for working with files
from os import makedirs
from os.path import join

# import the json library for writing out config files
import json

# import the subprocess library for running code from command line
import subprocess

# import the emlddmm image registration library
import sys
sys.path.append('../..')
import emlddmm
```

##### 7.2.2 Load images

[2]:
```
target_name = '/home/dtward/data/csh_data/emlddmm/mouse_example/NisslDown/'
atlas_name = '/home/dtward/data/AllenInstitute/allen_vtk/ara_nissl_50.vtk'
label_name = '/home/dtward/data/AllenInstitute/allen_vtk/annotation_50.vtk'
```

[3]:
```
# load the atlas with normalization (mean of abs is 1)
xI,I,_,_ = emlddmm.read_data(atlas_name,normalize=True)
# draw a picture
fig,ax = emlddmm.draw(I,xI,cmap='gray')
fig.suptitle('Atlas image')
```

```
<IPython.core.display.Javascript object>
```

```
<IPython.core.display.HTML object>
```

[3]:
```
Text(0.5, 0.98, 'Atlas image')
```

[4]:
```
# load the atlas segmentation labels, with no normalization (because these are integer␣
↪labels)
xS,S,_,_ = emlddmm.read_data(label_name)
SRGB = emlddmm.labels_to_rgb(S)
# draw a picture, showing labels and MRI
fig,ax = emlddmm.draw(SRGB+I/np.max(I)*2.0,xS)
fig.suptitle('Atlas image')
```

```
<IPython.core.display.Javascript object>
```

```
<IPython.core.display.HTML object>
```

[4]:
```
Text(0.5, 0.98, 'Atlas image')
```

[5]:
```
# load the target
xJ,J,_,_ = emlddmm.read_data(target_name)
# "weights" for missing data are stored in the last channel
W = J[-1]
J = J[:-1]
# draw a picture
fig,ax = emlddmm.draw(J,xJ,cmap='gray')
fig.suptitle('Target image image')
```

```
<IPython.core.display.Javascript object>
```

```
<IPython.core.display.HTML object>
```

```
<IPython.core.display.Javascript object>
```

```
<IPython.core.display.HTML object>
```

[5]:
```
Text(0.5, 0.98, 'Target image image')
```

##### 7.2.3 Perform an initial "slice to neighbor" alignment

[6]:
```
# downsample, to speed up calculations
xJd,Jd,Wd = emlddmm.downsample_image_domain(xJ,J,[1,2,2],W=W)
emlddmm.draw(Jd,xJd,vmin=0,vmax=1,interpolation='none')
```

```
<IPython.core.display.Javascript object>
```

```
<IPython.core.display.HTML object>
```

[6]:
```
(<Figure size 640x480 with 15 Axes>,
 array([[<AxesSubplot:>, <AxesSubplot:>, <AxesSubplot:>, <AxesSubplot:>,
         <AxesSubplot:>],
        [<AxesSubplot:>, <AxesSubplot:>, <AxesSubplot:>, <AxesSubplot:>,
         <AxesSubplot:>],
        [<AxesSubplot:>, <AxesSubplot:>, <AxesSubplot:>, <AxesSubplot:>,
         <AxesSubplot:>]], dtype=object))
```

[7]:
```
# run slice to neighbor alignment
# it gave terrible results!
# everything seemed to shift out of frame
import importlib
importlib.reload(emlddmm)
```

(continues on next page)

(continued from previous page)

```
out0 = emlddmm.atlas_free_reconstruction(xJ=xJd,J=Jd,W=(Wd==1),draw=True,n_steps=10,
↪eA2d=2e4)
```

```
../../emlddmm.py:4984: RuntimeWarning: divide by zero encountered in true_divide
  op = 1.0 / (xJ[0] - xJ[0][0])
../../emlddmm.py:4986: RuntimeWarning: divide by zero encountered in true_divide
  op = 1.0 / (xJ[0] - xJ[0][0])**2
../../emlddmm.py:4987: RuntimeWarning: divide by zero encountered in true_divide
  op = 1.0 / np.abs((xJ[0] - xJ[0][0]))**1.5/np.sign((xJ[0] - xJ[0][0]))
```

```
<IPython.core.display.Javascript object>
```

```
<IPython.core.display.HTML object>
```

```
<IPython.core.display.Javascript object>
```

```
<IPython.core.display.HTML object>
```

```
<IPython.core.display.Javascript object>
```

```
<IPython.core.display.HTML object>
```

```
<IPython.core.display.Javascript object>
```

```
<IPython.core.display.HTML object>
```

```
starting it 0
```

```
/home/dtward/.local/intelpython3/lib/python3.7/site-packages/torch/autograd/__init__.py:
↪199: UserWarning: grad and param do not obey the gradient layout contract. This is not␣
↪an error, but may impair performance.
grad.sizes() = [668, 3, 3], strides() = [9, 3, 1]
param.sizes() = [668, 3, 3], strides() = [9, 1, 3] (Triggered internally at ../torch/
↪csrc/autograd/functions/accumulate_grad.h:202.)
  allow_unreachable=True, accumulate_grad=True)  # Calls into the C++ engine to run the␣
↪backward pass
```

```
starting it 1
```

```
../../emlddmm.py:1404: UserWarning: To copy construct from a tensor, it is recommended␣
↪to use sourceTensor.clone().detach() or sourceTensor.clone().detach().requires_grad_
↪(True), rather than torch.tensor(sourceTensor).
  A2d = torch.tensor(A2d.detach().clone(),device=device, dtype=dtype)
```

```
starting it 2
starting it 3
starting it 4
starting it 5
starting it 6
starting it 7
starting it 8
starting it 9
```

##### 7.2.4 Run atlas to slice alignment

[8]:
```
config = {
    'device':'cpu', # cpu or cuda:0
    'downI':[[4,4,4],[2,2,2],[1,1,1]], # downsampling factors for the atlas for multi␣
↪scale
    'downJ':[[1,4,4],[1,2,2],[1,1,1]], # downsampling factors for the target for multi␣
↪scale. don't downsample in the slice direction (first number)
    'n_iter':[100,50,25], # number of iterations of gradient descent
    'a':[500.0], # spatial scale of the deformation
    'dv':[1000.0], # voxel size to sample the deformation on
    'muB':[[1.0,1.0,1.0]], # estimate of the intensity of the background (white)
    'muA':[[0.0,0.0,0.0]], # estimate of the intensitiy of artifacts (black)
    'slice_matching':[True], # enable rigid motions over slices
    'v_start':[0], # at which iteration do we start optimizing over the deformation
    'eA':[1e7], # gradient descent stepsize for 3D affine
    'eA2d':[1e5], # gradient descent stepsize for 2D rigid
    'ev':[1e-2], # gradient descent stepsize for deformation
    'local_contrast':[[1,16,16]], # divide the images into small blocks to estimate␣
↪contrast differences
    'up_vector':[[0.0,0.0,-1.0]], # what vector in the atlas should correspond to "up"␣
↪in a 2D image
    'sigmaR':[1e4] # regularization for deformation (bigger = less regularization)
}
# intial 2D alignment
config['A2d'] = out0['A2d']
# initial 3D affine
# look at the above figures and assin a letter to each row in order
# A means this row moves from posterior to anterior
# P means this row moves from anterior to posterior
# S means this row moves from inferior to superior
# I means this row moves from superior to inferior
# R means this row moves from left to right
# L means this row moves from right to left
# note that often it is difficult to tell left from right, so we always assume a right␣
↪handed coordinates
A = np.eye(4)
A[:3,:3] = emlddmm.orientation_to_orientation('ARI','PSL')
config['A'] = A
```

[9]:
```
import time
start = time.time()
import importlib
importlib.reload(emlddmm)
out = emlddmm.emlddmm_multiscale(xI=[xI],I=I,xJ=[xJ],J=J,W0=W,**config)
end = time.time()
print(end-start) # print out the elapsed time, which was reported in the manuscript
```

```
Found 3 scales
```

```
<IPython.core.display.Javascript object>
```

```
<IPython.core.display.HTML object>
```

```
<IPython.core.display.Javascript object>
```

```
<IPython.core.display.HTML object>
```

```
<IPython.core.display.Javascript object>
```

```
<IPython.core.display.HTML object>
```

```
<IPython.core.display.Javascript object>
```

```
<IPython.core.display.HTML object>
```

```
<IPython.core.display.Javascript object>
```

```
<IPython.core.display.HTML object>
```

```
<IPython.core.display.Javascript object>
```

```
<IPython.core.display.HTML object>
```

```
<IPython.core.display.Javascript object>
```

```
<IPython.core.display.HTML object>
```

```
<IPython.core.display.Javascript object>
```

```
<IPython.core.display.HTML object>
```

```
<IPython.core.display.Javascript object>
```

```
<IPython.core.display.HTML object>
```

```
../../emlddmm.py:176: RuntimeWarning: invalid value encountered in true_divide
  J /= (vmax[:,None,None,None] - vmin[:,None,None,None])
```

```
Iteration 70, linear oscilating, reducing eA to 9000000.0
Iteration 80, linear oscilating, reducing eA to 8100000.0
```

```
../../emlddmm.py:1335: UserWarning: To copy construct from a tensor, it is recommended␣
↪to use sourceTensor.clone().detach() or sourceTensor.clone().detach().requires_grad_
↪(True), rather than torch.tensor(sourceTensor).
  v = torch.tensor(v.detach().clone(),device=device,dtype=dtype)
../../emlddmm.py:1379: UserWarning: To copy construct from a tensor, it is recommended␣
↪to use sourceTensor.clone().detach() or sourceTensor.clone().detach().requires_grad_
↪(True), rather than torch.tensor(sourceTensor).
  A = torch.tensor(A.detach().clone(),device=device,dtype=dtype)
```

```
<IPython.core.display.Javascript object>
```

```
<IPython.core.display.HTML object>
```

```
<IPython.core.display.Javascript object>
```

```
<IPython.core.display.HTML object>
```

```
../../emlddmm.py:1449: RuntimeWarning: More than 20 figures have been opened. Figures␣
↪created through the pyplot interface (`matplotlib.pyplot.figure`) are retained until␣
↪explicitly closed and may consume too much memory. (To control this warning, see the␣
↪rcParam `figure.max_open_warning`).
  figA2d,axA2d = plt.subplots(2,2)
```

```
<IPython.core.display.Javascript object>
```

```
<IPython.core.display.HTML object>
```

```
<IPython.core.display.Javascript object>
```

```
<IPython.core.display.HTML object>
```

```
<IPython.core.display.Javascript object>
```

```
<IPython.core.display.HTML object>
```

```
<IPython.core.display.Javascript object>
```

```
<IPython.core.display.HTML object>
```

```
<IPython.core.display.Javascript object>
```

```
<IPython.core.display.HTML object>
```

```
<IPython.core.display.Javascript object>
```

```
<IPython.core.display.HTML object>
```

```
<IPython.core.display.Javascript object>
```

```
<IPython.core.display.HTML object>
```

```
Iteration 20, linear oscilating, reducing eA to 9000000.0
Iteration 40, linear oscilating, reducing eA to 8100000.0
```

```
<IPython.core.display.Javascript object>
```

```
<IPython.core.display.HTML object>
```

```
<IPython.core.display.Javascript object>
```

```
<IPython.core.display.HTML object>
```

```
<IPython.core.display.Javascript object>
```

```
<IPython.core.display.HTML object>
```

```
<IPython.core.display.Javascript object>
```

```
<IPython.core.display.HTML object>
```

```
<IPython.core.display.Javascript object>
```

```
<IPython.core.display.HTML object>
```

```
<IPython.core.display.Javascript object>
```

```
<IPython.core.display.HTML object>
```

```
<IPython.core.display.Javascript object>
```

```
<IPython.core.display.HTML object>
```

```
<IPython.core.display.Javascript object>
```

```
<IPython.core.display.HTML object>
```

```
<IPython.core.display.Javascript object>
```

```
<IPython.core.display.HTML object>
```

```
715.610100030899
```

##### 7.2.5 Apply the transform from atlas to registered space

We use the inverse transform.

[10]:
```
tform = emlddmm.compose_sequence(
    [
        emlddmm.Transform(out[-1]['A'],direction='b'),
        emlddmm.Transform(out[-1]['v'],domain=out[-1]['xv'],direction='b'),
    ],
    xJ
)
AphiI = emlddmm.apply_transform_float(xI,I,tform)
AphiS = emlddmm.apply_transform_int(xS,S,tform)
```

##### 7.2.6 Apply the transform from target to registered space

We use the forward transform

[11]:
```
tform = emlddmm.compose_sequence(
    [
        emlddmm.Transform(out[-1]['A2d'],direction='f'),
    ],
    xJ
)
RiJ = emlddmm.apply_transform_float(xJ,J,tform)
```

[12]:
```
# draw the labels over the target image
AphiSRGB = emlddmm.labels_to_rgb(AphiS,white_label=0)
# draw a picture, showing labels and MRI
fig,ax = emlddmm.draw(AphiSRGB*0.125+RiJ.numpy()/RiJ.max().numpy()*0.875,xJ)
fig.suptitle('Target image with atlas labels')
```

```
<IPython.core.display.Javascript object>
```

```
<IPython.core.display.HTML object>
```

[12]:
```
Text(0.5, 0.98, 'Target image with atlas labels')
```

##### 7.2.7 Apply the transform from registered target to atlas space

Interpolation between slices is only meaningful after 2D alignment. Here we apply a transformation to our aligned Nissl slices.

[24]:
```
tform = emlddmm.compose_sequence(
    [
        emlddmm.Transform(out[-1]['v'],domain=out[-1]['xv'],direction='f'),
        emlddmm.Transform(out[-1]['A'],direction='f'),
    ],
    xI
)
phiiAiRiJ = emlddmm.apply_transform_float(xJ,RiJ,tform)
```

[27]:
```
fig,ax = emlddmm.draw(phiiAiRiJ)
```

```
<IPython.core.display.Javascript object>
```

```
<IPython.core.display.HTML object>
```

##### 7.2.8 Run this example from the command line

We will write out the parameters above to a config file, and run our command line interface. This will produce all our standard outputs.

##### 7.2.9 Write out the registration config file

We'll make sure some of the config inputs are formatted properly as plain text for our config file, and write it out..

[13]:
```
config_ = dict(config)
config_['A2d'] = [config_['A2d'].tolist()]
config_['A'] = [config_['A'].tolist()]
```

[14]:
```
config_file = 'atlas_to_target_config.json'
with open(config_file,'wt') as f:
    json.dump(config_,f)
```

##### 7.2.10 Write out the transformation graph config file

[15]:
```
transformation_config = {
  "output": "output",
  "space_image_path": [
    [
      "Atlas",
      "image",
      atlas_name
    ],
    [
      "Atlas",
      "labels",
      label_name
    ],
    [
      "Target",
      "image",
      target_name
    ],
  ],
  "registrations": [
    [
      [
        "Atlas",
        "image"
      ],
```

(continues on next page)

(continued from previous page)

```
      [
        "Target",
        "image"
      ]
    ],
  ],
  "configs": [
    config_file,

  ],
  "transform_all":True,
}
transformation_config_file = 'transformation_graph_config.json'
with open(transformation_config_file,'wt') as f:
    json.dump(transformation_config,f)
```

##### 7.2.11 Run the command

[16]:
```
command = f'python -u ../../transformation_graph_v01.py --infile {transformation_config_
↪file} > outputs.txt 2>&1'
print('about to run command:')
print(command)
```

```
about to run command:
python -u ../../transformation_graph_v01.py --infile transformation_graph_config.json >␣
↪outputs.txt 2>&1
```

[17]:
```
subprocess.call(command,shell=True)
```

[17]:
```
0
```

##### 7.2.12 View all the outputs

All the output directories are printed below. Note that this includes a python file graph.p, which contains information about the transformation graph. This will allow us to apply the transforms we have already calculated to new datasets later.

[20]:
```
from os import walk
for dirpath,dirnames,filenames in walk('output',):
    for i,f in enumerate(filenames):
        if i >= 3:
            print('...more files for each slice...')
            break
        print(join(dirpath,f))
```

```
output/infile.json
output/graph.p
output/Target_registered/Target_to_Target_registered/transforms/Target_registered_PTM902-
↪N1-2021.05.27-15.39.29_PTM902_3_0001_to_Target_PTM902-N1-2021.05.27-15.39.29_PTM902_3_
↪0001_matrix.txt
```

(continues on next page)

(continued from previous page)

```
output/Target_registered/Target_to_Target_registered/transforms/Target_registered_PTM902-
↪N1-2021.05.27-15.39.29_PTM902_2_0002_to_Target_PTM902-N1-2021.05.27-15.39.29_PTM902_2_
↪0002_matrix.txt
output/Target_registered/Target_to_Target_registered/transforms/Target_registered_PTM902-
↪N1-2021.05.27-15.39.29_PTM902_1_0003_to_Target_PTM902-N1-2021.05.27-15.39.29_PTM902_1_
↪0003_matrix.txt
...more files for each slice...
output/Target_registered/Target_to_Target_registered/images/Target_image_PTM902-N1-2021.
↪05.27-15.39.29_PTM902_3_0001_to_Target_registered_PTM902-N1-2021.05.27-15.39.29_PTM902_
↪3_0001.vtk
output/Target_registered/Target_to_Target_registered/images/Target_image_PTM902-N1-2021.
↪05.27-15.39.29_PTM902_2_0002_to_Target_registered_PTM902-N1-2021.05.27-15.39.29_PTM902_
↪2_0002.vtk
output/Target_registered/Target_to_Target_registered/images/Target_image_PTM902-N1-2021.
↪05.27-15.39.29_PTM902_1_0003_to_Target_registered_PTM902-N1-2021.05.27-15.39.29_PTM902_
↪1_0003.vtk
...more files for each slice...
output/Target_registered/Atlas_to_Target_registered/qc/Target_image_registered.jpg
output/Target_registered/Atlas_to_Target_registered/qc/Atlas_image_to_Target_registered.
↪jpg
output/Target_registered/Atlas_to_Target_registered/images/Atlas_image_to_Target_
↪registered_PTM902-N1-2021.05.27-15.39.29_PTM902_3_0001.vtk
output/Target_registered/Atlas_to_Target_registered/images/Atlas_image_to_Target_
↪registered_PTM902-N1-2021.05.27-15.39.29_PTM902_2_0002.vtk
output/Target_registered/Atlas_to_Target_registered/images/Atlas_image_to_Target_
↪registered_PTM902-N1-2021.05.27-15.39.29_PTM902_1_0003.vtk
...more files for each slice...
output/Target_registered/Atlas_to_Target_registered/transforms/Target_registered_PTM902-
↪N1-2021.05.27-15.39.29_PTM902_3_0001_to_Atlas_displacement.vtk
output/Target_registered/Atlas_to_Target_registered/transforms/Target_registered_PTM902-
↪N1-2021.05.27-15.39.29_PTM902_2_0002_to_Atlas_displacement.vtk
output/Target_registered/Atlas_to_Target_registered/transforms/Target_registered_PTM902-
↪N1-2021.05.27-15.39.29_PTM902_1_0003_to_Atlas_displacement.vtk
...more files for each slice...
output/Atlas/Target_registered_to_Atlas/transforms/velocity.vtk
output/Atlas/Target_registered_to_Atlas/transforms/A.txt
output/Atlas/Target_registered_to_Atlas/transforms/Atlas_to_Target_registered_
↪displacement.vtk
output/Atlas/Target_registered_to_Atlas/qc/Target_image_to_Atlas.jpg
output/Atlas/Target_registered_to_Atlas/qc/Atlas_image.jpg
output/Atlas/Target_to_Atlas/images/Target_image_to_Atlas.vtk
output/Atlas/Target_to_Atlas/transforms/Atlas_to_Target_displacement.vtk
output/Target/Atlas_to_Target/qc/Atlas_image_to_Target.jpg
output/Target/Atlas_to_Target/qc/Target_image.jpg
output/Target/Atlas_to_Target/images/Atlas_image_to_Target_PTM902-N1-2021.05.27-15.39.29_
↪PTM902_3_0001.vtk
output/Target/Atlas_to_Target/images/Atlas_image_to_Target_PTM902-N1-2021.05.27-15.39.29_
↪PTM902_2_0002.vtk
output/Target/Atlas_to_Target/images/Atlas_image_to_Target_PTM902-N1-2021.05.27-15.39.29_
↪PTM902_1_0003.vtk
...more files for each slice...
output/Target/Atlas_to_Target/transforms/Target_PTM902-N1-2021.05.27-15.39.29_PTM902_3_
```

(continues on next page)

(continued from previous page)

```
↪0001_to_Atlas_displacement.vtk
output/Target/Atlas_to_Target/transforms/Target_PTM902-N1-2021.05.27-15.39.29_PTM902_2_
↪0002_to_Atlas_displacement.vtk
output/Target/Atlas_to_Target/transforms/Target_PTM902-N1-2021.05.27-15.39.29_PTM902_1_
↪0003_to_Atlas_displacement.vtk
...more files for each slice...
output/Target/Target_registered_to_Target/transforms/Target_PTM902-N1-2021.05.27-15.39.
↪29_PTM902_3_0001_to_Target_registered_PTM902-N1-2021.05.27-15.39.29_PTM902_3_0001_
↪matrix.txt
output/Target/Target_registered_to_Target/transforms/Target_PTM902-N1-2021.05.27-15.39.
↪29_PTM902_2_0002_to_Target_registered_PTM902-N1-2021.05.27-15.39.29_PTM902_2_0002_
↪matrix.txt
output/Target/Target_registered_to_Target/transforms/Target_PTM902-N1-2021.05.27-15.39.
↪29_PTM902_1_0003_to_Target_registered_PTM902-N1-2021.05.27-15.39.29_PTM902_1_0003_
↪matrix.txt
...more files for each slice...
```

CHAPTER

### EIGHT

### FUNCTION REFERENCE

CHAPTER

### NINE

### WORK IN PROGRESS

This section contains a description of works in progress.

#### 9.1 Blockface imaging

Our pipeline supports a sequence of blockface photographs for 3D reconstruction rather than a tranditional 3D volume. We will prepare examples related to this application.

CHAPTER

TEN

### INSTALLING

- Make sure you have python 3 installed with pip
- Use pip to install the packages in the requirements.txt file (pip in stall -r requrements.txt)
- Clone the repository on github (git clone https://github.com/twardlab/emlddmm)
- When running interactively in python, make sure you add the path (import sys; sys.path.append('/LOCATION/OF/REPOSITORY'))

CHAPTER

### ELEVEN

### EXAMPLES

We include data and code for two examples, in the "examples" folder. Both examples show code run interactively in a jupyter notebook, and show how the command line interface is used.

- 3D Human MRI example
- Mouse Nissl serial section alignment example.

CHAPTER

TWELVE

### IMPORTANT FUNCTIONS

#### 12.1 Python functions to be run interactively

- `emlddmm.emlddmm()`: Run the emlddmm algorithm.
- `emlddmm.emlddmm_multiscale()`: Run the emlddmm algorithm iteratively at different scales.

#### 12.2 Command line functions

- `transformation_graph_v01`: Run registration between two or more datasets using the transformation graph command line interface.

CHAPTER

THIRTEEN

### MODULE AND FUNCTION DOCUMENTATION

All functions are automatically documented with sphinx and napoleon. See the Function Reference section.

CHAPTER

### FOURTEEN

### WEB INTERFACE

To improve accessibility, we provide a web interface at https://twardlab.com/reg which can be used for small jobs. Please email the Daniel Tward to request an account. We provide a guest account for review purposes. Username: guest, password: 84983c60. No identifying information is recorded in the guest account. We request email addresses when creating your own account.
