## Supplementary material for "Solving the *where* problem and quantifying geometric variation in neuroanatomy using generative diffeomorphic mapping": workflow

### Registration workflow supplement

Supplementary material for Tward, et al. 2024

#### Introduction

In this supplement we summarize the steps that a user undertakes to register serial sections of a mouse brain to an annotated reference atlas.

#### Procedure

**Data preparation:** A brain id is provided by users and the location of the corresponding images is determined. A basic quality check is first performed automatically, including checking section number, image size, data structure, etc. If quality is as expected, and data ready to be registered, the high resolution images will be downsampled by a factor of 32 by 32, saved in TIF format, and sent to the registration pipeline.

**Registration:** The core of our registration code is written in python. It will first determine the modality of the brain images. Our pipeline is currently being applied to sections alternately imaged with Nissl and fluorescence signals, or Nissl and immunohistochemistry (IHC) staining. A simple initial slice alignment of Nissl images is applied by translating the center of mass of each section to the center of the image, followed by a rigid alignment of slices to a weighted combination of their nearest neighbors. An initial affine transformation is computed between the reference atlas and this initial stack, followed by deformable registration using the procedure outlined in the manuscript. Nissl images are rigidly registered to their adjacent images of the alternate modality. Transformation parameters (deformation fields, linear transformation matrices, scale change information) are output in VTK format (see the data format supplementary material), and several images are produced for quality control.

**Registration quality control and correction**: An online viewer was built for registration quality control (QC) purposes. During the registration step, a low resolution image with atlas overlay is generated for each section and these images are used for QC. An example of our QC interface is shown in Figure 1.

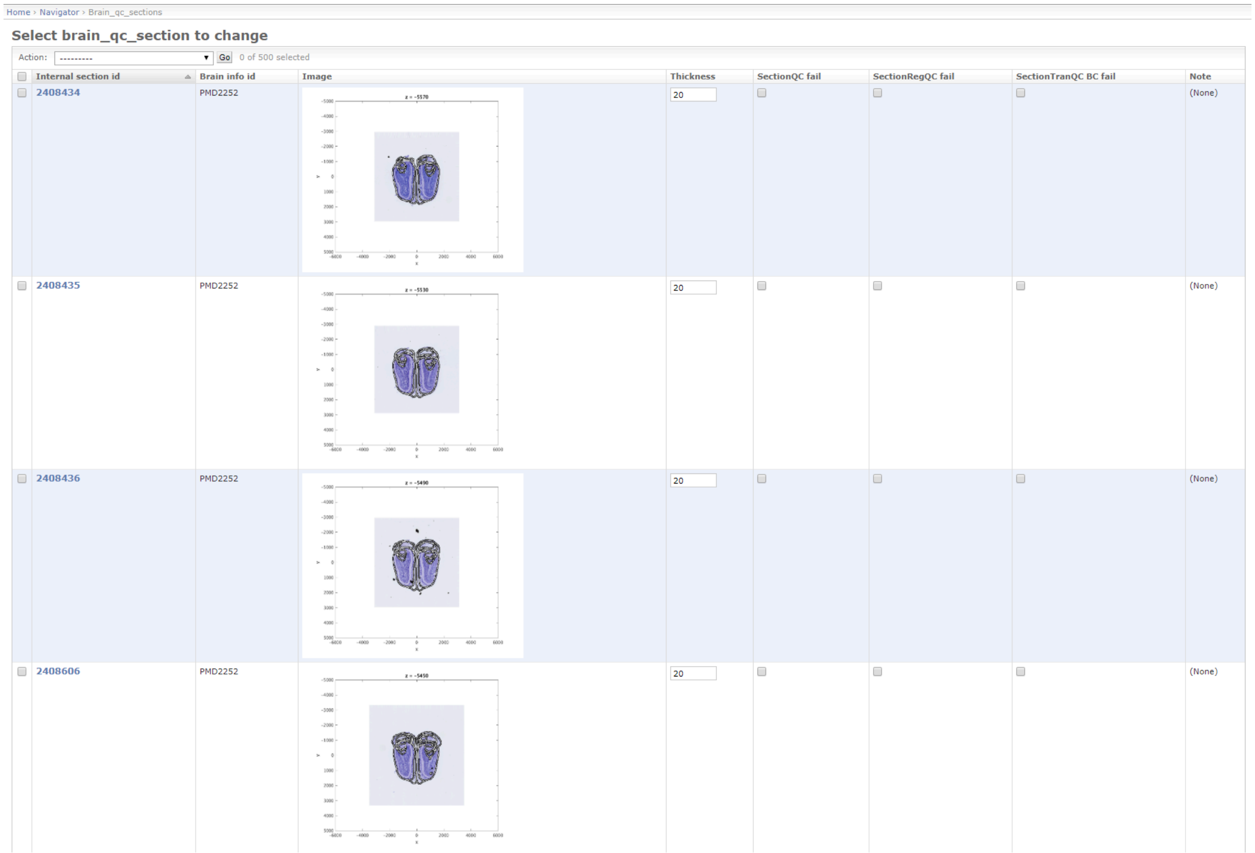


**Figure 1:** Web based quality control interface allows users to view registered sections with atlas annotation overlays and verify imaging parameters before the expensive process of transforming high resolution images begins.

Accurate rigid alignment between Nissl sections and sections of other modalities is critical, and these mappings occasionally fail due to findinging local minima in a nonconvex optimization problem. We have created an interactive tool where a user chooses corresponding landmark points in pairs of images. A rigid transformation is displaced and updated with each new landmark. This manual alignment can either be used directly, or treated as an initial guess for the automatic rigid alignment optimization procedure.

**Transformation:** The high resolution images can be transformed after the registered brain passes QC or is manually corrected. Due to the quality of the tape transfer method, only rigid registration is applied to each 2D brain section, which can be applied very quickly using matrix arithmetic, as opposed to interpolating a displacement vector field. Each section is transformed and padded to a fixed image size of 24000 * 24000 pixels. The 10um atlas is deformed nonlinearly and upsampled to image resolution. In order to display the image and annotation on the viewer, brain sections are compressed into JP2 format, and annotation is converted into geojson.

**Running time:** Registration is run on a shared supercomputer cluster at Cold Spring Harbor Laboratory (CSHL). Since it is a shared resource, the number of jobs that can run at the same time varies. We require about 16 CPU threads and 24G memory for one brain to finish in 8 hours. Compute time varies considerably depending on data and parameters chosen however, and the serial section alignment example we include in our github repository takes only about 12 minutes to complete (for 668 Nissl slices). Typically, 30 brains can be registered in parallel. Applying the transformation to high resolution images requires relatively high memory, and therefore is run on a multi-node custom built cluster. The cluster has 8 nodes and each node has 72 CPU threads, 188G memory and 2 Nvidia GTX 2080TI GPUs. Each node can transform one brain in 2.5 hours, so the maximum capacity is about 80 brains per day.
