## Supplementary material for "Solving the *where* problem and quantifying geometric variation in neuroanatomy using generative diffeomorphic mapping": diffeomorphometry

### Diffeomorphometry supplement

Supplementary material for Tward, et al. 2024

---

#### Introduction

Understanding typicality and variability in brain anatomy as a quantitative science has been the goal of the Computational Anatomy program[1]. Several important mathematical and algorithmic results have been established, which we draw upon for accurate brain registration and statistical analysis. The key feature of this field is to model observed anatomical images as spatial deformations of a well characterized template. Employing the deformable template approach allows these transformations, known as diffeomorphisms, to become the objects of study when describing shape, rather than images themselves. The quantitative study of shape using diffeomorphisms has become known as diffeomorphometry [2]. In this supplement we review some of these important details.

#### Diffeomorphisms and their action on images

Diffeomorphic transformations are differentiable invertible coordinate mappings with a differentiable inverse. Given a background space $X$ (typically a subset of $\mathbb{R}^2$ or $\mathbb{R}^3$ real numbers), we denote them by $\varphi : X \to X$, which maps $x \mapsto \varphi(x)$. The set of diffeomorphisms form a mathematical group under the composition operation ($[\varphi_A \circ \varphi_B](x) = \varphi_A(\varphi_B(x))$), with identity elements, existence of inverse, and associativity (but not commutativity). This set is not a vector space, meaning linear combinations of diffeomorphisms does not generally lead to a diffeomorphism, making standard linear modeling techniques inappropriate.

Given an anatomical image $I : X \to \mathbb{R}$, diffeomorphisms deform them via composition with the inverse $\varphi \cdot I = I \circ \varphi^{-1}$. While sometimes counterintuitive, this left action obeys necessary associativity properties $\varphi_B \cdot (\varphi_A \cdot I) = (\varphi_B \circ \varphi_A) \cdot I$. It is immediately clear that working with invertible transformations is essential to transform points ($x_i \mapsto \varphi(x_i)$) and images ($I \mapsto I \circ \varphi^{-1}$) consistently.

#### Derivatives define scale change

Because $\varphi$ is a diffeomorphism, its Jacobian matrix $D\varphi$ is well defined and invertible everywhere in space. This matrix defines how a small cube in the atlas is transformed into a small paralelipiped in the observed volume. The ratio of the paralellipiped's volume to that of the cube is determined by the determinant of the Jacobian. Working with the cube root changes units from volume to length, and we call the result the scale change. Studying this object as a means to quantify local size and shape differences has been well established [3].

#### Generating diffeomorphisms via smooth flows

In computational anatomy diffeomorphisms are generated by integrating time varying velocity fields, which can be thought of colloquially as “displacement equals velocity times time”. We denote a time varying velocity field $v_t : X \to \mathbb{R}^D$ where $D$ is typically 2 or 3. A diffeomorphism is constructed by integrating Euler’s equation from time 0 to 1

$$\frac{d}{dt}\varphi_t = v_t(\varphi_t),\ \text{with}\ \varphi_0 = \text{identity}\ . \qquad (1)$$

The inverse can be constructed from the optical flow equation:

$$\frac{d}{dt}\varphi_t^{-1} = -D\varphi_t^{-1}v_t,\ \text{with}\ \varphi_0^{-1} = \text{identity}\ . \qquad (2)$$

In image registration we work with the inverse equation which is solved numerically using Semi Lagrangian advection[4]. In our notation, when no time index is specified on $\varphi_t$ time $t = 1$ is implied.

#### The vector space of smooth flows

To guarantee that integrating a velocity field leads to a diffeomorphism, it must have a sufficient number of continuous derivatives [5]. This is achieved in practice by modeling $v_t$ as belonging to a Reproducing Hilbert Space of smooth functions $V$, with an inner product that penalizes high frequency components:

$$\langle u, v\rangle_V = \int [Lu]^T(x)[Lv](x)dx, \quad L = (\text{identity} - \alpha^2\text{Laplacian})^2 \qquad (3)$$

where $\alpha$ is a parameter with units of length that controls the length scale of smoothness. This leads to an associated norm which allows us to speak of length and distance in the space of diffeomorphisms, $\|v\|_V^2 = \langle v, v\rangle_V$.

#### Registration as an optimal control problem

Image registration can be posed as an optimization problem that balances accuracy in a least squares sense with smoothness in terms of the norm defined above. An atlas image $I$ can be transformed to a target image $J$ by solving the optimization problem:

$$v^* = \arg\max_v \frac{1}{2\sigma_R^2} \int_0^1 \|v_t\|_V^2 dt + \frac{1}{2\sigma_M^2} \int |I(\varphi_1^{-1}(x)) - J(x)|^2 dx \ . \qquad (4)$$

For this problem, $\alpha$, $\sigma_R^2$, and $\sigma_M^2$ are user specified constants that define the balance between smoothness scale, smoothness strength, and matching accuracy (respectively). This problem was originally solved in [6], and has been approached by various other researchers (and ourselves) with different accuracy functionals.

#### Optimal trajectories are geodesics

An important property of optimal solutions is that $v_t^*$ defines a length minimizing curve between identity and $\varphi_1$. These geodesic curves are characterized by a conservation of momentum equation [7]. Let $m = L^* L v$ . Here $*$ refers to the adjoint of the linear operator, which is defined implicitly by

$$\int [Lu]^T(x)[Lv](x)dx = \int [u]^T(x)[L^* Lv](x)dx \ . \qquad (5)$$

Note that the operator $L$ defined above is self adjoint ($L^* = L$). The geodesic curves are characterized by

$$\frac{d}{dt} m + Dmv + m\text{div}v + Dv^T m = 0 \ . \qquad (6)$$

Therefore, given a velocity vector field defined at $t = 0$, one can reconstruct the entire optimal trajectory $v_t$ and thus the diffeomorphism $\varphi_1$. Since $v_0$ lies in the vector space $V$, we use $v_0$ as a linear parameterization of $\varphi_1$. In the language of if differential geometry, $v_0$ is a vector in the tangent space to the diffeomorphism group at the identity element.

#### Tangent space kernel PCA

Because $v_0$ lies in a linear space, it can be modeled (when sampled on a voxel grid) as a multivariate Gaussian. In this setting we can use principal component analysis to compute uncorrelated, orthogonal modes of variability. Because we have defined $v_0$ as belonging to the vector space $V$, orthogonal must be interpreted as $\langle u, v \rangle_V = 0$. This procedure is described in detail in [8], and we summarize it here.

We let $N$ be the number of voxels in our atlas image, and $M$ be the number of samples of $v_0$ we have measured. Let $X$ be a $3N \times M$ data matrix formed by vectorizing and stacking each $v_0$, and $X_0$ be centered by subtracting the mean at each voxel $\bar{X}$. Instead of working with the covariance matrix directly, we compute the Gram matrix of inner products as $G = X_0^T K X_0$ where $K$ is the $3N \times 3N$ kernel matrix that applies $L^* L$ to our vectorized velocity fields. In practice $K$ is applied using Fourier transforms, and not stored as a matrix. We then compute the eigendecomposition $G = VDV^T$, for

$V$ unitary and $D$ diagonal. The variance of the $i$-th orthogonal mode is given by $\sigma_i^2 = D_{ii}/N$, and the modes themselves are given by the columns of $U = X_0 V D^{-\frac{1}{2}}$, which are orthogonal with respect to our inner product.

For the $i$-th mode of variation, we can initialize Eq (6) with $v_0 = \bar{X} + s\sigma_i U_{\cdot i}$ and reconstruct a diffeomorphism $\varphi_1$ that is $s$ standard deviations from the mean in the direction of the first mode. From this diffeomorphism we can analyze scale change as described above.

#### References


1. Grenander, U. & Miller, M. I. Computational anatomy: An emerging discipline. *Quart. Appl. Math.* **56**, 617–694 (1998).
2. Miller, M. I., Younes, L. & Trouvé, A. Diffeomorphometry and geodesic positioning systems for human anatomy. *Technology* **2**, 36 (2014).
3. Ashburner, J. & Friston, K. J. Voxel-based morphometry—the methods. *Neuroimage* **11**, 805–821 (2000).
4. Staniforth, A. & Côté, J. Semi-Lagrangian Integration Schemes for Atmospheric Models—A Review. *Mon. Weather Rev.* **119**, 2206–2223 (1991).
5. Dupuis, P., Grenander, U. & Miller, M. I. VARIATIONAL PROBLEMS ON FLOWS OF DIFFEOMORPHISMS FOR IMAGE MATCHING. *Quart. Appl. Math.* **56**, 587–600 (1998).
6. Beg, M. F., Miller, M. I., Trouvé, A. & Younes, L. Computing Large Deformation Metric Mappings via Geodesic Flows of Diffeomorphisms. *Int. J. Comput. Vis.* **61**, 139–157 (2005).
7. Miller, M. I., Trouvé, A. & Younes, L. Geodesic Shooting for Computational Anatomy. *J. Math. Imaging Vis.* **24**, 209–228 (2006).
8. Vaillant, M., Miller, M. I., Younes, L. & Trouvé, A. Statistics on diffeomorphisms via tangent

space representations. *Neuroimage* **23 Suppl 1**, S161–9 (2004).
