## Supplementary material for "Solving the *where* problem and quantifying geometric variation in neuroanatomy using generative diffeomorphic mapping": stereology

### Sterology correction factor

Supplementary material for Tward, et al. 2024

---

#### Introduction

In this supplement we summarize how we derived our stereological correction factor for cell density analysis.

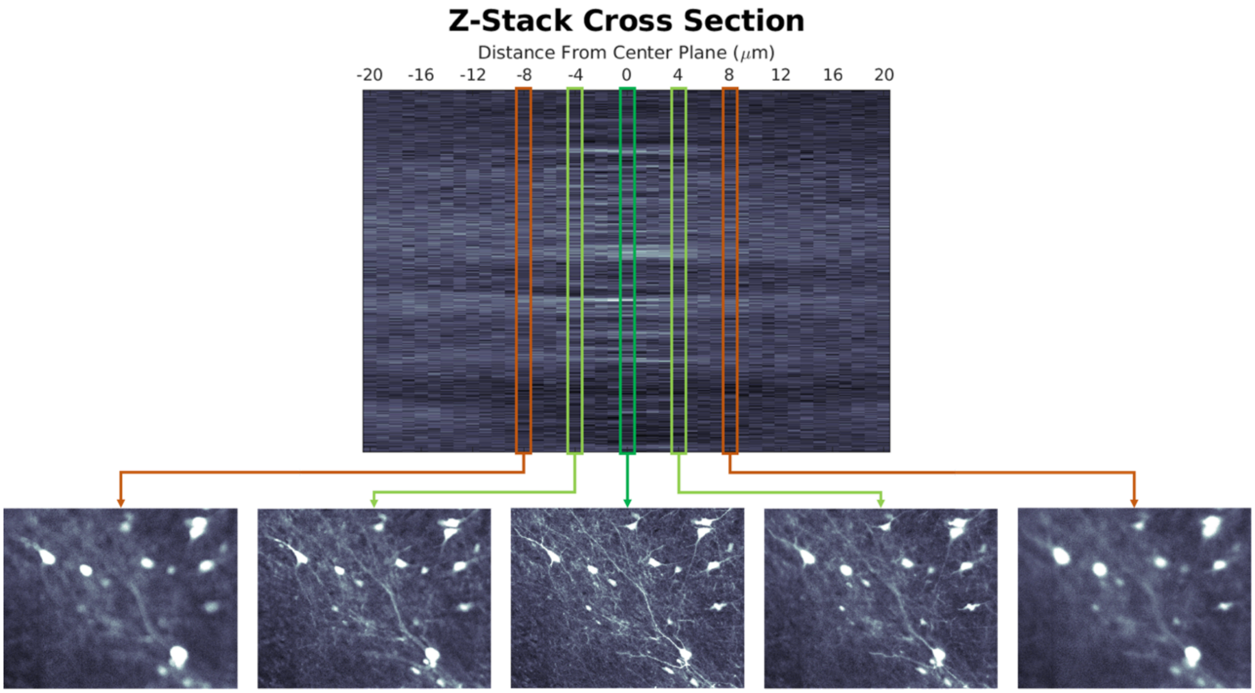


Figure S.1. Top panel: a cross section of a window in a z-stack obtained using the same imaging system as analyzed datasets. Note the bright stripes of signal; each corresponds to a neuron or cluster of adjacent neurons. Bottom panel: Example images of the window in the z-stack at the center plane of the section and also at 4μm and 8μm from the center plane in both directions. Green boxes indicate all cell bodies and processes are still in focus (at 4μm) while orange boxes (at 8μm) indicate loss of focus leading to loss of defined cell bodies or processes.

#### Stereological correction procedure

We calculate an effective post-processing section thickness and attendant cell density estimation correction factor based on a z-stack obtained on the same imaging system as the datasets analyzed in the present manuscript (Nanozoomer HT 2.0). The top panel Figure S.X shows a cross-section of a window through the z-stack of section cut with the microtome set for 20µm section thickness, with distance from the center plane in the z-direction given in µm. The estimation procedure is detailed below:

1. We let the original section thickness be T (which is set during microtomy to 20µm).
    a. Note that cutting on the microtome is subjected to some variation (1-2 µm) around the nominal mean value of 20µm. Cutting will produce both slightly thicker and thinner sections. To a first approximation we therefore adopt a constant section thickness of 20µm
2. Let the measured in-plane density (from running cell detection) be $\rho_{2D}$. We need to know the relation between $\rho_{2D}$ and $\rho_{3D}$, where the latter is the 3D volumetric density. The relation between the two may be written as:
    a. $\rho_{3D} = \rho_{2D} / t$, where t = an effective section thickness for converting the 2D densities to 3D densities.
3. The data shows that the histological processing shrinks the physical thickness of the section. We estimate the effective post-processing section thickness, to be 10-12µm as follows:
    a. In example images (bottom panel of Figure S.1) of the focal plane at 4µm from the center plane in either direction, we note clear neuronal cell bodies and processes in focus. By 8µm from the center plane, the imaged z-plane is out of focus; however, cell bodies are still discernible, suggesting this z-plane is outside the edge of the tissue in both directions.
    b. In the top panel of Figure S.1 showing the z-direction cross section, we note that the brightest fluorescent signals (each one indicates a cell or group of adjacent cells) go to ~5-6µm from the center plane, lending support to a post processing section thickness between 10-12µm.
4. Based on these observations we assume that if we detect every object in the optical section, small and large, we will capture any cell that is wholly or partially in that section. With this assumption (which we justify based on the z-stack data above), the effective thickness of a slab over which the observed cell centers are distributed, is given by t=T+2R, where R is the radius of the cell nuclei (which we assume to be spheres for this purpose; note that the in-plane profiles we observe are largely circular).
5. Assuming that the cell centers follow a spatial Poisson distribution with a density parameter of $\rho_{3D}$, we then have the following equation:
    a. $\rho_{2D} = \rho_{3D} \times (T+2d)$, or $\rho_{3D} = \rho_{2D} / (T+2R)$

Setting T=20µm and R=5µm we have an effective thickness of 30µm. We use this conversion factor to compute the 3D densities from the 2D densities in the original image data (before any diffeomorphic mapping).
