## Supplementary material for "Solving the *where* problem and quantifying geometric variation in neuroanatomy using generative diffeomorphic mapping": refinement

### Mapping refinement

Supplementary material for Tward, et al. 2024

#### Introduction

Quality control and mapping accuracy is key for scientific insight. While annotations that result from atlas mapping can be updated by expert anatomists, our analysis relies on a consistency between annotations and mappings that are also studied. We therefore refine mappings based on manual annotations as opposed to updated annotations directly.

#### The manual annotation cost function

If the result of a mapping is not satisfactory, users may annotate observed images $J^i$ with one or more binary masks $M^i$, each corresponding to some set of labels in Allen Reference Atlas. A synthesized version $\hat{M}^i$ is generated from the Allen atlas, which is expected to match only in the region indicated (i.e. the user need only perform segmentations in regions with inaccuracies, not everywhere in the image).

$$\arg \min_{\varphi^0, A, \varphi^i, R^i, f^i} \mathrm{Reg}^0(\varphi^0) + \sum_i \mathrm{Reg}^i(\varphi^i) + \frac{1}{2\sigma^2} \int |\hat{J}^i(x) - J^i(x)|^2 \pi^i(x) dx + \frac{1}{2\sigma_M^2} \int |\hat{M}^i(x) - M^i(x)|^2 M^i(x) dx \ . \qquad (1)$$

This minimization problem is initialized using the previous solution, which is typically close to optimal, reducing computation time needed.
